# DeSpotX: Identifiability-Based Decontamination for Spatial Transcriptomics

**DOI:** 10.64898/2026.05.12.724704

**Authors:** Ruo Han Wang, Andrew J. Gentles

## Abstract

Spatial transcriptomics (ST) at single-cell resolution profiles gene expression in its native spatial context, but a substantial fraction of transcripts contaminate neighboring cells, compromising downstream biological analyses. Existing decontamination methods rely on heuristic priors and either ignore the spatial structure of contamination or aggregate over neighbors without separating contamination from native expression, leaving the decomposition ambiguous. To resolve this ambiguity, we introduce DeSpotX, a deep generative model that uses anchor genes, defined as genes not natively expressed in a given cell cluster, to constrain the contamination decomposition and make it identifiable. DeSpotX further uses spatial information to estimate contamination locally through a cluster-masked, distance-weighted average over neighboring cells, and prevents over-correction of low-expression signal through a learned diffusion prior. On spike-in simulations across five datasets and four ST platforms, DeSpotX achieves AUROC *>* 0.94 on every dataset, with gains of 0.02 to 0.12 over the best baseline, and remains robust to inaccuracies in the cell-cluster annotation and in anchor gene construction. On real tissues, we show that the decontaminated counts produce improved marker-gene specificity, more spatially coherent expression, and cell-cell communication networks consistent with known biology. We further show that iterating decontamination and cell-cluster annotation refines these outcomes, reassigning ligand-receptor signaling to the expected source cells in mouse brain and breast cancer tissues.

## 1 Introduction

Single-cell-resolution spatial transcriptomics (ST) platforms, such as Xenium [1], MERFISH [2, 3], CosMx [4], and Stereo-seq [5], now profile millions of cells in tissue, enabling the study of cellular organization, cell-cell communication, and disease microenvironments at single-cell resolution [6, 7]. However, these technologies suffer from a systematic contamination artifact, where transcripts from one cell are frequently assigned to neighboring cells through ambient diffusion, segmentation errors, and vertical overlap of cells in the tissue slice [8, 9, 10]. The contamination is widespread, with 20-40% of cells showing substantial contamination and 10-20% of cells expressing markers of mutually exclusive cell types according to recent studies [8, 9]. These artifacts compromise downstream biological analyses, causing misannotation of cell types, blurring the spatial organization of gene expression, and producing spurious cell-cell communication networks that can mislead the discovery of therapeutic targets.

Decontamination of ST data poses three fundamental challenges. First, the native and contamination components cannot be uniquely separated from observed counts. The same observed count distribution can arise from multiple combinations of native expression, contamination profile, and contamination fraction, so methods fail to identify which combination is correct without additional constraints on the decomposition [11]. Second, contamination in ST is spatially structured, since it diffuses locally from nearby cells and the contaminating signal reflects the cellular composition of each cell’s local environment. A single global ambient profile, as used by methods designed for dissociated single-cell RNA-seq [12, 13, 14], fails to capture this local structure. Third, low-expression genes are vulnerable to over-correction during contamination removal [15], even though they often carry informative biological signal. At low expression levels, even small errors in the contamination estimate are large relative to the true signal, so the subtraction can remove real expression along with contamination.

To address each of these challenges, we present DeSpotX (https://github.com/Gentles-lab/DeSpotX), an identifiability-based deep generative model for decontaminating single-cell-resolution ST data. For the *non-identifiability* challenge, DeSpotX introduces anchor genes, defined as genes that are not natively expressed in a given cell cluster and inferred automatically from per-cluster expression rates, and uses them to constrain the decomposition. For the *spatial structure* challenge, DeSpotX estimates the local contamination profile as a cluster-masked, distance-weighted average over cross-cluster spatial neighbors, separating spatially diffused contamination from each cell’s native expression. For the *signal preservation* challenge, DeSpotX places a learned diffusion prior over the latent state encoding native expression, regularizing recovered profiles toward learned cluster-conditioned distributions and preventing the noise in contamination estimates from over-correcting low-expression signal.

Across five spatial transcriptomics datasets spanning four platforms and four tissue contexts, DeSpotX outperforms all leading decontamination baselines and maintains strong performance even when cell-cluster annotations or anchor gene construction are imperfect. Downstream analyses on the decontaminated counts yield cleaner cluster separation, more spatially coherent marker expression, and more biologically coherent cell-cell communication networks. We further show that iterating between decontamination and cell-cluster annotation progressively improves these outcomes.

## 2 Related work

### Decontamination of single-cell RNA-seq

Several methods correct ambient RNA contamination in dissociated single-cell RNA-seq. SoupX [13] estimates a global ambient profile from empty droplets and subtracts a per-cell scaled version. DecontX [12] fits a Bayesian mixture model that decomposes each cell’s expression into its native cluster profile and a global contamination profile aggregated across clusters, recovering per-cell contamination fractions by variational inference. CellBender [14] models raw barcode-level counts with a learned ambient prior estimated from empty droplets, separating ambient transcripts from cell-derived expression. These methods rely on a single global ambient profile and do not incorporate spatial information, making them unable to capture the spatial structure of contamination in single-cell-resolution ST.

### Decontamination of spatial transcriptomics

Recent methods adapt count correction to ST data by incorporating spatial information into the contamination estimate. SpaceBender [10] replaces CellBender’s global ambient profile with a spatially local profile computed by averaging expression over *k*-nearest-neighbor or distance-based spatial neighbors. DenoIST [9] fits a Poisson mixture that classifies each gene count as endogenous or contamination based on the cell’s count relative to its local neighborhood. ResolVI [8] decomposes observed counts into native expression, neighbor-derived diffusion, and non-specific background. While these methods leverage spatial structure, none separate spatially diffused contamination from native expression at each location. Moreover, all of these methods regularize the decomposition through Bayesian priors, latent-variable architectures, or mixture-model assumptions, none of which resolve the underlying non-identifiability. These regularizers can produce decompositions that fit the observed counts, but the recovered decompositions can deviate from the true profile. A detailed comparison between DeSpotX and existing methods is provided in Appendix A.

### Identifiability of latent variable models

Identifiability is a long-standing concern in latent variable models, where multiple parameter combinations can produce the same observed data distribution [11]. Two main strategies resolve non-identifiability. The first uses auxiliary variables to make latent dimensions conditionally independent, as in identifiable VAEs [11], but no such auxiliary variable is available in the decontamination setting. The second imposes structural constraints on the model, such as anchor-word identifiability in topic models [16] and separability in non-negative matrix factorization [17], and fits the decontamination problem naturally, since each cell cluster expresses only a subset of genes. DeSpotX builds on this strategy, proving that the contamination decomposition is non-identifiable from observed counts alone and deriving an anchor-gene constraint that provably restores identifiability. To our knowledge, DeSpotX is the first decontamination method for spatial transcriptomics derived from a formal identifiability analysis, replacing prior-based regularization with a provable guarantee.

## 3 Methods

### 3.1 Problem formulation

Consider a single-cell-resolution spatial transcriptomics dataset of *N* cells indexed by *i* ∈ {1, …, *N* }. For each cell *i*, we observe a count vector 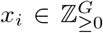 over *G* genes, a discrete cluster label *t*_*i*_ ∈ {1, …, *T*}, and a spatial coordinate *s*_*i*_ ∈ ℝ^2^ . Let 𝒩 (*i*) ⊂ {1, …, *N*} with |𝒩 (*i*)| = *K* denote the *K* nearest spatial neighbors of cell *i* under the Euclidean metric. We model the observed counts as a mixture of the cell’s native transcriptional output and contamination from its local spatial environment:

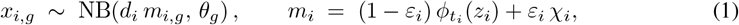

where *d*_*i*_ = Σ_*g*_ *x*_*i,g*_ is the library size, *θ*_*g*_ ∈ ℝ_*>*0_ is a gene-wise dispersion parameter, 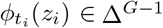 is the cell’s native expression profile over genes conditioned on its cluster index *t*_*i*_ and a latent state 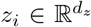, *χ*_*i*_ ∈ Δ^*G−*1^ is the local contamination profile at location *s*_*i*_, and *ε*_*i*_ ∈ [0, 1] is the per-cell contamination fraction. We adopt the mean-dispersion parameterization throughout, where NB(*µ, θ*) denotes the negative binomial with mean *µ* and variance *µ* + *µ*^2^*/θ*.

The inference task is to recover, for each cell, the native profile 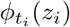, the local contamination profile *χ*_*i*_, and the contamination fraction *ε*_*i*_ from the observation *x*_*i*_, the cluster index *t*_*i*_, and the spatial context {(*x*_*j*_, *t*_*j*_, ∥*s*_*j*_ − *s*_*i*_ ∥) : *j* ∈ 𝒩 (*i*)} . The likelihood in Eq. (1) depends on (*ε*_*i*_, 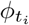, *χ*_*i*_) only through the mixture *m*_*i*_, making the decomposition non-identifiable.

#### Lemma 1

(Non-identifiability of the native–contamination decomposition). *For any observed mean m* ∈ Δ^*G−*1^, *the set of parameters consistent with m*,

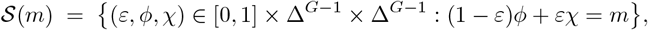

*contains a continuum of solutions on which the likelihood in* *Eq*. (1) *is constant*.

A proof is provided in Appendix B.1. Consequently, any decontamination method must impose external constraints on at least one of *ϕ, χ*, or *ε* to recover a unique decomposition. DeSpotX imposes such constraints through anchor positions in *ϕ* and a spatial estimator for *χ*, and mitigates noise sensitivity through a latent diffusion prior over *z*.

### 3.2 DeSpotX architecture

DeSpotX instantiates Eq. (1) as a deep generative model with four components (Figure 1). A **spatial graph encoder** maps each cell’s spatial context to a latent encoding *z*_*i*_ of its native expression state and an estimate of the contamination fraction 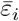, alongside a local contamination profile *χ*_*i*_ aggregated from cross-cluster spatial neighbors. A **latent diffusion prior** constrains *z*_*i*_ to a data-driven distribution of plausible cluster-conditioned latents, preserving biological signal under noisy contamination estimates. A **cluster-conditioned decoder** maps (*z*_*i*_, *t*_*i*_) to the native profile 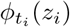. **Anchor constraints**, inferred automatically from per-cluster expression rates in the data, restore identifiability of the decomposition by fixing *ϕ*_*t,g*_ to zero at (*t, g*) pairs where gene *g* is not natively expressed in cluster *t*.

**Figure 1:**
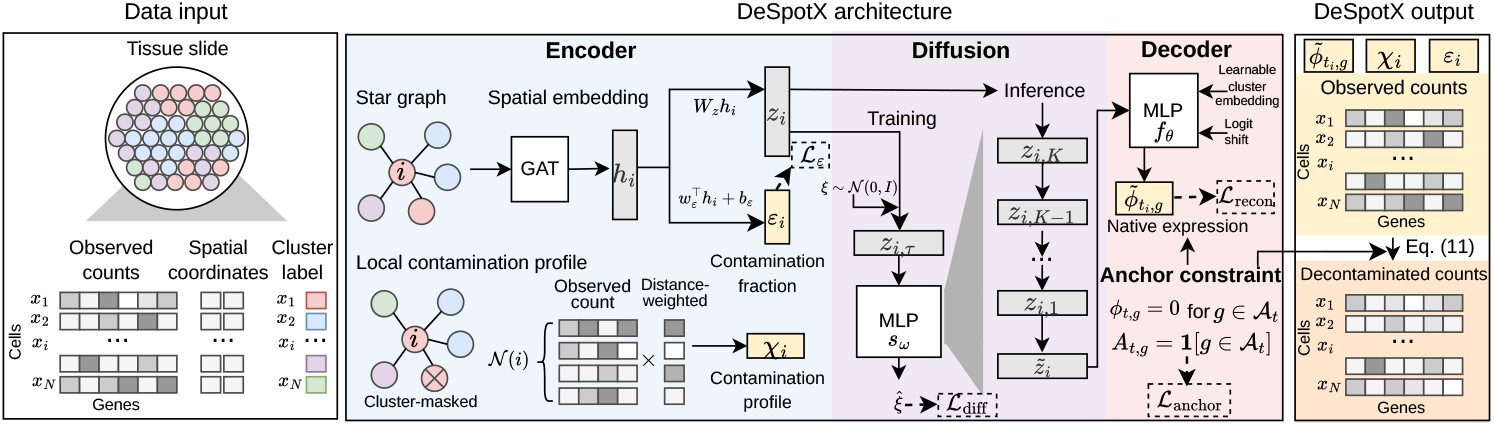
Overview of DeSpotX for decontaminating spatial transcriptomics data.

#### Spatial graph encoder

For each cell *i* we construct a star graph 𝒢_*i*_ in which a single center node, cell *i*, is connected to *K* leaf nodes corresponding to the cells in 𝒩 (*i*). Node features are log-transformed counts concatenated with a cluster embedding; edge features are kernel weights *w*_*ij*_ = exp(− ∥*s*_*j*_ − *s*_*i*_∥ */ρ*_*i*_) with adaptive bandwidth *ρ*_*i*_ set to the median distance from cell *i* to its *K* nearest neighbors. A GATv2 layer [18] aggregates neighbor information into the center, producing a spatial-context embedding 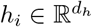. From *h*_*i*_, the encoder produces a latent state *z*_*i*_ and a contamination-fraction estimate 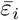:

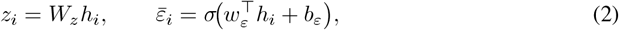

where *W*_*z*_, *w*_*ε*_, and *b*_*ε*_ are learned parameters, and the sigmoid ensures 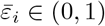. Implementation details are provided in Appendix C.1.

We estimate the local contamination profile *χ*_*i*_ as a cluster-masked, distance-weighted average over 𝒩 (*i*):

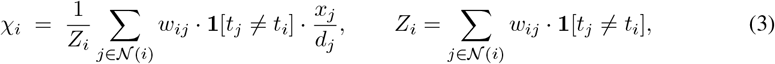

with fallback to a dataset-level cross-cluster ambient profile when no other-cluster neighbor is available. The mask **1**[*t*_*j*_ ≠ *t*_*i*_] restricts *χ*_*i*_ to a cross-cluster aggregation of the local spatial environment, isolating shared contamination from the center cell’s own native signal. This masking excludes within-cluster contamination; the resulting error in 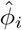 is bounded by 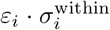, where 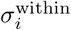 measures within-cluster expression heterogeneity (Appendix B.2, validated in Appendix I).

#### Latent diffusion prior

The spatial estimator *χ*_*i*_ carries error that propagates into the recovered decomposition. Writing 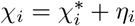 with 𝔼[*η*_*i*_] = 0, solving the mixture equation for 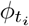 gives

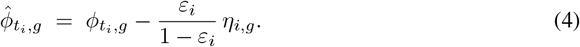

The relative distortion 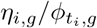 grows large when 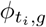 is small, leading to spurious removal of low-but-real signal. We address this by regularizing *z* with a diffusion prior over plausible cluster-conditioned latents.

We model the distribution of cluster-conditioned latents via a denoising diffusion process [19]. A score network *s*_*ω*_ is trained alongside the encoder and decoder to predict noise added to encoder latents:

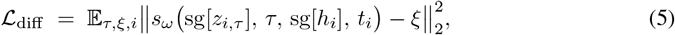

where *z*_*i,τ*_ is the noised latent at diffusion step *τ, ξ* is the injected noise, and sg[·] denotes stop-gradient, which prevents the encoder from collapsing *z* to a degenerate distribution that *s*_*ω*_ trivially denoises. At inference, *z*_*i*_ is refined by a DDIM [20] reverse trajectory, projecting it onto the learned manifold before decoding. Implementation details are provided in Appendix C.2.

#### Cluster-conditioned decoder

The decoder maps the latent state and cluster index to the native expression profile:

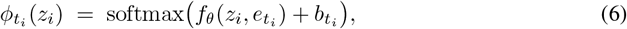

where *f*_*θ*_ is a multilayer perceptron, 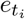 is a learned cluster embedding, and 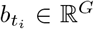 is a cluster-specific logit shift initialized from the log-mean expression of cluster *t*_*i*_. Implementation details are provided in Appendix C.3.

#### Anchor constraints

By Lemma 1, identifiability requires external constraints. We introduce *anchor genes*: for each cluster *t*, let 𝒜_*t*_ ⊆ {1, …, *G*} denote a set of genes not natively expressed in cluster *t*, and let *A*_*t,g*_ = **1**[*g* ∈ 𝒜_*t*_]. The set 𝒜_*t*_ is identified from per-cluster expression rates (Appendix C.4).

##### Lemma 2

(Identifiability of the decomposition under anchor constraints). *Fix a cell i with cluster index t*_*i*_, *and assume χ*_*i*_ *is constructed without using x*_*i*_. *Suppose ϕ*_*t,g*_ = 0 *for all g* ∈ 𝒜_*t*_, *and that χ*_*i,g*_ *>* 0 *for at least one* 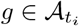. *Then ε*_*i*_ *is identifiable from the observations at anchor positions, and* 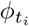 *is identifiable on non-anchor positions whenever ε*_*i*_ *<* 1.

A proof is provided in Appendix B.3.

#### Training objective

The four components are trained together under

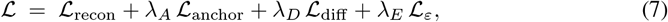

where the reconstruction loss is the per-cell negative log-likelihood under Eq. (1),

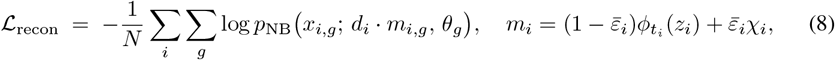

the anchor penalty

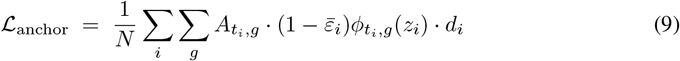

penalizes nonzero 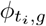 at anchor positions, the diffusion loss _diff_ is given in Eq. (5), and the contamination-fraction regularizer

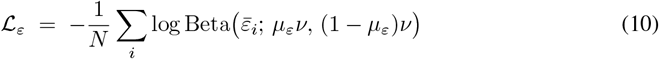

pulls 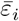 toward a learnable target *µ*_*ε*_ with fixed concentration *v*. The anchor weight *λ*_*A*_ is linearly increased from zero over the first *E*_warm_ epochs, letting the decoder establish reasonable profiles before the constraint is enforced fully. Optimization details are provided in Appendix C.5.

### 3.3 Inference

With all components fixed after training, denoised counts are produced as follows. The star graph encoder yields the spatial-context embedding *h*_*i*_, the latent state *z*_*i*_, the contamination fraction 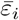, and the local contamination profile *χ*_*i*_. DDIM refinement then projects *z*_*i*_ onto the learned manifold, and the resulting 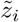 is decoded into the refined native profile 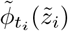. Each observation is then reweighted by the native component’s contribution to the mixture:

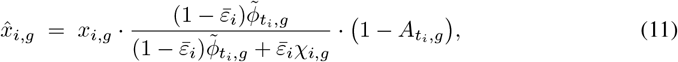

where the final factor sets anchor positions to zero in the output.

## 4 Experiments

### 4.1 Datasets

We evaluate DeSpotX and baseline methods on five publicly available single-cell-resolution spatial transcriptomics datasets (Table 1). The five datasets span the major imaging- and sequencing-based platforms (Xenium [1], Xenium 5K Prime [21], CosMx [4], MERFISH [3], and Stereo-seq [5]), four tissue contexts (human breast cancer, human non-small cell lung cancer, mouse brain, and mouse embryo), and panel sizes ranging from 313 to 18,582 genes. Cell-cluster annotations are taken from the original studies where available; for Xenium5k_Breast, annotations are obtained by CellTypist [22] label transfer using a published human breast atlas as reference [23]. All datasets are used as published, without additional quality control or filtering; raw integer counts are provided to all methods, with spatial centroids and cell-cluster annotations additionally provided to methods that use them.

**Table 1:** Single-cell-resolution spatial transcriptomics datasets.

| Dataset name | Technology | Tissue | # cells | # genes | # cell clusters |
| --- | --- | --- | --- | --- | --- |
| Xenium_Breast [1] | Xenium | Breast cancer | 118,370 | 313 | 20 |
| MERFISH_Brain [3] | MERFISH | Mouse brain | 115,779 | 550 | 29 |
| CosMx_NSCLC [4] | CosMx | Non-small cell lung cancer | 99,185 | 980 | 38 |
| Xenium5k_Breast [21] | Xenium 5K Prime | Breast cancer | 577,258 | 5,101 | 7 |
| Stereo-seq_E14 [5] | Stereo-seq | Mouse embryo (E14.5) | 92,928 | 18,582 | 16 |

### 4.2 Spike-in simulations

Spatial transcriptomics data lack ground truth for ambient contamination, so we construct a spike-in benchmark from real datasets.

#### 4.2.1 Simulation design

To prevent the benchmark from favoring DeSpotX, we sample contamination counts from a Poisson likelihood (rather than DeSpotX’s negative-binomial) and draw per-cell contamination rates from a Beta distribution (rather than DeSpotX’s regularized encoder estimate). For each of the five datasets in Table 1, we sample 40,000 cells and inject synthetic ambient counts at six noise levels 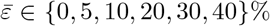. For each cell, we (i) construct a local ambient profile from its nearest cross-cell-cluster neighbors, (ii) draw a per-cell contamination rate from a Beta distribution with mean 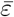, and (iii) sample per-gene contamination counts from the resulting per-cell, per-gene Poisson rates. The spiked counts *s*_*i,g*_ = *n*_*i,g*_ + *c*_*i,g*_ are provided to each method as input, while the native counts *n*_*i,g*_, the per-gene contamination labels 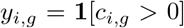, and the realized contamination fraction *ε*_true_ = Σ_*i,g*_ *c*_*i,g*_*/* Σ_*i,g*_ *s*_*i,g*_ are retained as ground truth. Details are provided in Appendix D.

#### 4.2.2 Benchmark evaluation

#### Baselines

We compare DeSpotX against four published decontamination baselines, including SoupX [13], DecontX [12], ResolVI [8], and SpaceBender [10], all run with default settings. Cell-Bender [14] is excluded because it requires empty-droplet observations that are unavailable for the ST platforms, and DenoIST [9] is excluded because it requires per-molecule transcript files that are unavailable in our benchmark. All methods receive the same spiked counts *s*_*i,g*_ as input; spatial coordinates and cell-cluster labels are additionally provided to methods that use them. Each method outputs decontaminated counts 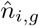.

#### Metrics

We evaluate each method on three complementary metrics. (i) AUROC scores how well each method’s removed contamination 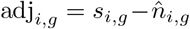 predicts the binary label *y*_*i,g*_ = **1**[*c*_*i,g*_ *>* 0] of synthetic contamination. (ii) per-cell calibration error (PCE) and (iii) global calibration error (GCE) measure the absolute deviation between each method’s implied contamination fraction and the true rate, computed per cell and dataset-wide respectively. Since spatial transcriptomics data contain platform-derived contamination, both PCE and GCE are computed after subtracting each method’s removal at 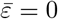.

#### Benchmark performance

Table 2 reports each metric averaged across the five non-zero noise levels, with DeSpotX achieving the best performance on every metric and every dataset. On AUROC, DeSpotX is uniformly above 0.94, while the next-best method, ResolVI, averages 0.892 across the five datasets. The non-spatial methods, SoupX and DecontX, achieve AUROC scores below 0.74 on every dataset, indicating that contamination in ST data is spatially structured. DeSpotX also achieves the lowest PCE and GCE on every dataset, with the methods ranking similarly on PCE and GCE, indicating that DeSpotX’s global calibration arises from correct per-cell localization rather than averaging out per-cell errors. DeSpotX’s advantage is widest on the largest gene panels (Xenium5k_Breast and Stereo-seq_E14), where its GCE is 1.32 and 3.72 percentage points (pp) respectively, while no other method falls below 8 pp on Xenium5k_Breast or 13 pp on Stereo-seq_E14.

**Table 2:** Benchmark comparison across five spatial transcriptomics datasets. We report AUROC, per-cell calibration error (PCE, in percentage points), and global calibration error (GCE, in percentage points). Values are mean ± std across the non-zero noise levels. Best value per row in bold.

| Dataset | Metric | DeSpotX | SoupX [13] | DecontX [12] | ResolVI [8] | SpaceBender [10] |
| --- | --- | --- | --- | --- | --- | --- |
| Xenium_Breast | AUROC↑ | **0.955 ± 0.007** | 0.606 ± 0.069 | 0.677 ± 0.014 | 0.851 ± 0.002 | 0.805 ± 0.009 |
| | PCE↓ | **15.79 ± 10.39** | 16.45 ± 9.87 | 18.35 ± 12.48 | 20.62 ± 14.27 | 16.33 ± 9.58 |
| | GCE↓ | **10.03 ± 4.78** | 11.26 ± 4.60 | 13.11 ± 6.84 | 15.71 ± 8.76 | 10.21 ± 6.04 |
| MERFISH_Brain | AUROC↑ | **0.947 ± 0.008** | 0.631 ± 0.008 | 0.721 ± 0.013 | 0.812 ± 0.009 | 0.824 ± 0.013 |
| | PCE↓ | **10.83 ± 7.14** | 20.66 ± 13.94 | 16.99 ± 11.75 | 17.60 ± 13.58 | 12.72 ± 7.79 |
| | GCE↓ | **5.27 ± 1.92** | 16.08 ± 8.41 | 12.97 ± 6.87 | 12.19 ± 7.98 | 7.57 ± 3.68 |
| CosMx_NSCLC | AUROC↑ | **0.967 ± 0.003** | 0.696 ± 0.011 | 0.683 ± 0.002 | 0.886 ± 0.005 | 0.700 ± 0.013 |
| | PCE↓ | **13.64 ± 9.22** | 15.43 ± 11.55 | 20.13 ± 13.45 | 21.15 ± 12.76 | 16.09 ± 9.87 |
| | GCE↓ | **7.64 ± 3.90** | 9.31 ± 6.37 | 14.52 ± 7.57 | 15.61 ± 7.34 | 12.61 ± 6.08 |
| Xenium5k_Breast | AUROC↑ | **0.997 ± 0.000** | 0.705 ± 0.024 | 0.732 ± 0.006 | 0.980 ± 0.000 | 0.604 ± 0.020 |
| | PCE↓ | **8.06 ± 4.39** | 16.16 ± 10.61 | 21.74 ± 14.30 | 19.07 ± 13.79 | 13.54 ± 7.38 |
| | GCE↓ | **1.32 ± 1.15** | 8.09 ± 5.16 | 15.57 ± 8.20 | 13.15 ± 7.94 | 10.74 ± 5.16 |
| Stereo-seq_E14 | AUROC↑ | **0.984 ± 0.001** | 0.719 ± 0.051 | 0.721 ± 0.002 | 0.929 ± 0.004 | 0.904 ± 0.010 |
| | PCE↓ | **15.74 ± 8.71** | 22.49 ± 17.03 | 19.78 ± 13.58 | 21.12 ± 13.91 | 18.86 ± 11.81 |
| | GCE↓ | **3.72 ± 1.93** | 17.87 ± 11.44 | 15.11 ± 8.05 | 16.51 ± 8.36 | 13.71 ± 6.54 |

#### Ablation study

We assess each architectural component by removing it from the full model on Xenium_Breast and MERFISH_Brain (full results in Appendix E). Removing the identifiability constraints produces the largest degradation, with GCE rising from 10 to over 70 pp on Xenium_Breast and from 5 to 17 pp on MERFISH_Brain. Removing the spatial graph encoder, the cluster mask, or the diffusion prior each reduces performance on all three metrics, confirming that every component contributes to DeSpotX’s accuracy. Beyond aggregate metrics, the diffusion prior specifically protects low-expression marker genes from over-correction (full results in Appendix F).

#### Robustness and consistency

DeSpotX is robust to the construction of the anchor mask, the perturbation of the cell-cluster annotation, the number of spatial neighbors *K*, and the cell-cluster annotation method (Appendix G). The performance of DeSpotX is consistent across contamination levels, and its margin over baselines grows at higher contamination levels (Appendix H). The within-cluster contamination bound is validated on spike-in simulations (Appendix I).

### 4.3 Application to real spatial transcriptomics data

We apply DeSpotX to the full five datasets in Table 1; it runs much faster than other ST methods (Appendix J). We then assess whether decontamination improves downstream analyses.

#### 4.3.1 Cell-cluster identity

We first examine whether decontamination produces clearer separation of cell clusters and tighter localization of marker-gene expression to canonical clusters. Figure 2a shows UMAP embeddings of the raw counts and the decontaminated counts from DeSpotX and the four baselines across the five datasets. Each panel reports the *k*NN purity at *k* = 20, defined as the fraction of each cell’s *k* nearest neighbors in UMAP space that share its cluster label; higher values indicate cleaner cluster separation. DeSpotX yields the highest *k*NN purity on every dataset and produces visibly better-separated clusters than the raw counts, while the baselines yield embeddings with less clear cluster separation.

**Figure 2:**
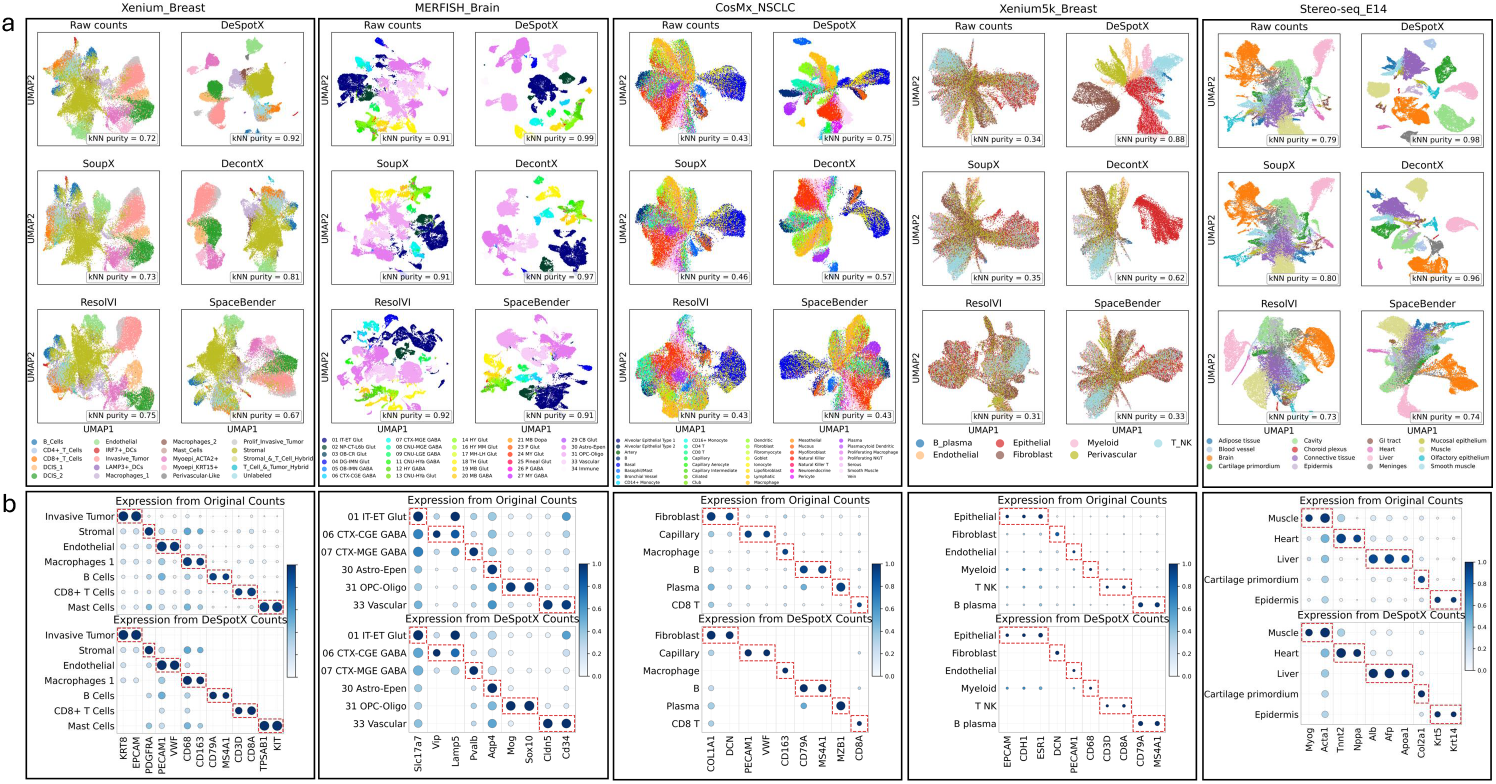
DeSpotX yields better-separated cluster structure and improved marker-gene specificity on real spatial transcriptomics data. (a) UMAP embeddings of raw counts and decontaminated counts from each method, across the five datasets, colored by cell cluster. *k*NN purity is shown in each panel. (b) Marker-gene dotplots for raw and DeSpotX-decontaminated counts across cell clusters in each dataset; red boxes mark canonical marker-cluster pairs.

Figure 2b compares marker-gene dotplots for the raw counts and the DeSpotX-decontaminated counts across cell clusters in each dataset. After decontamination, expression of canonical markers is reduced in non-expressing clusters while remaining concentrated in their expected clusters (red boxes). This pattern is consistent with the removal of ambient contamination from non-expressing cells, with native expression preserved in expressing cells. The same comparison for the four baseline methods is provided in Appendix K.

#### 4.3.2 Spatial coherence

We next assess whether DeSpotX decontamination preserves the spatial structure of marker expression. Figure 3a-e shows the spatial distribution of canonical cell-cluster markers in the raw and DeSpotX-decontaminated counts. In the raw counts, marker expression is detected throughout the tissue, including in regions where the canonical expressing cluster is absent. After DeSpotX, marker expression is concentrated within the canonical clusters and the surrounding tissue is largely cleared, indicating that DeSpotX removes ambient signal in non-expressing regions while preserving expression in regions where the marker is natively transcribed. Spatial maps of the removed counts further show that DeSpotX’s removal is spatially structured, concentrating near canonical expressing clusters, consistent with the assumption that contamination spreads locally from source cells (Appendix L).

**Figure 3:**
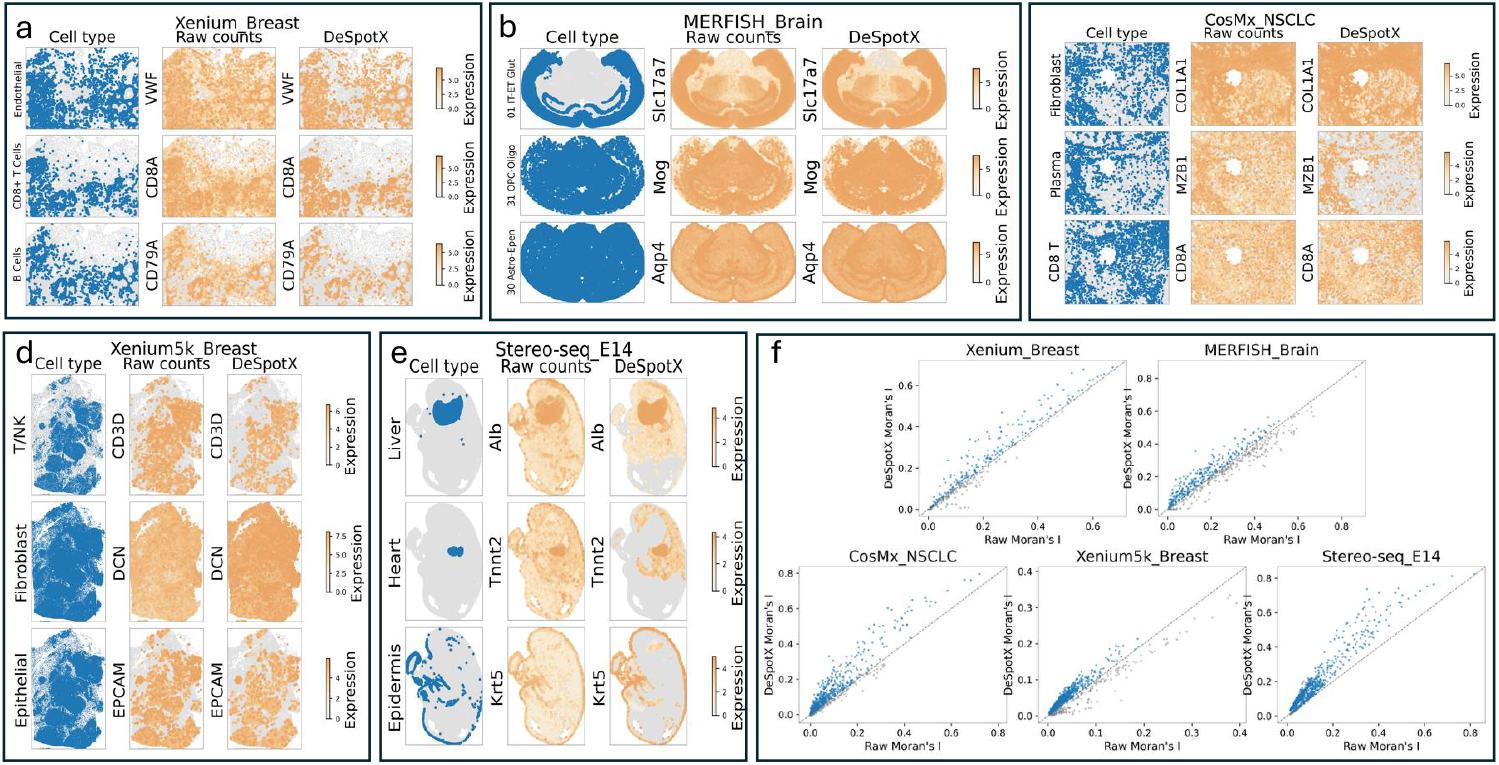
DeSpotX-decontaminated expression is spatially coherent. (a-e) Spatial maps of canonical cell-cluster markers in each dataset, comparing raw and DeSpotX-decontaminated counts. (f) Pergene Moran’s *I* for raw and DeSpotX-decontaminated counts across the five datasets.

Figure 3f reports per-gene Moran’s *I* for the raw and DeSpotX-decontaminated counts across the five datasets. Moran’s *I* measures whether gene expression is spatially autocorrelated, with higher values indicating that cells with similar expression are spatially close. Most genes show higher Moran’s *I* after DeSpotX, with the largest gains in genes that already exhibit some spatial structure in the raw data, indicating that decontamination preserves and strengthens biologically meaningful spatial patterns. The same comparison for baseline methods is provided in Appendix M.

#### 4.3.3 Iterative decontamination and cell-cluster annotation

DeSpotX requires a cell-cluster annotation as input, but the quality of this annotation depends on the data, which itself contains contamination. We therefore evaluate whether iterating between decontamination and cell-cluster annotation produces progressively better results. At each iteration, DeSpotX decontaminates the raw counts using the previous iteration’s annotation, and the decontaminated counts are re-annotated by a CellTypist classifier [22] trained on a reference atlas [24] (Figure 4a).

**Figure 4:**
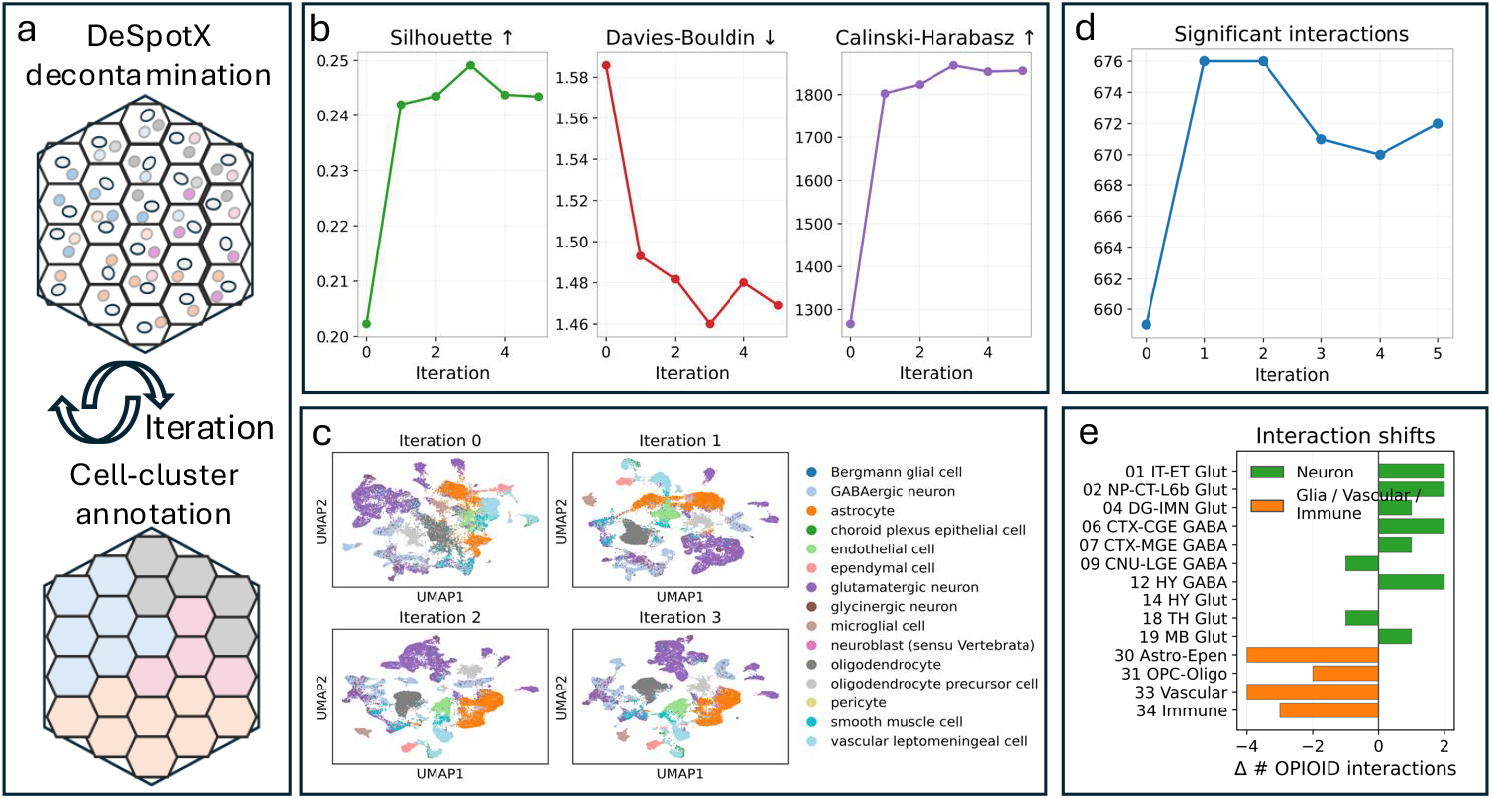
Iterative DeSpotX decontamination and cell-cluster annotation on `MERFISH_Brain`. (a) Schematic of the iterative procedure. (b) Cluster metrics across iterations. (c) UMAP embeddings of the decontaminated counts at iterations 0-3. (d) Total number of significant cell-cell interactions inferred by CellChat across iterations. (e) Change in the number of significant OPIOID-pathway interactions per sender cell-cluster between iteration 0 and iteration 5.

We apply this procedure to `MERFISH_Brain` for five iterations. Cluster-validity metrics improve and stabilize by iteration 3 (Figure 4b), with Silhouette rising from 0.20 to 0.24, Davies-Bouldin falling from 1.59 to 1.47, and Calinski-Harabasz increasing from 1,267 to 1,856. The UMAP embeddings show progressively cleaner separation of the brain cell types (Figure 4c). We run CellChat [25] on inferred ligand-receptor signaling networks at each iteration. The number of significant interactions stabilizes by iteration 2-3 (Figure 4d). Within the OPIOID pathway, the sender distribution shifts (Figure 4e), moving away from glial, vascular, and immune senders toward neuronal populations, consistent with the neuronal origin of opioid peptides [26, 24]. The same iterative procedure applied to `Xenium_Breast` also shows improvements in downstream analyses (Appendix N).

## 5 Discussion

We have presented DeSpotX, the first decontamination method for spatial transcriptomics derived from a formal identifiability analysis. Whereas existing methods rely on heuristic priors, we prove that the contamination decomposition is non-identifiable from observed counts alone and that anchor genes provably restore identifiability. DeSpotX outperforms existing methods on every benchmarked dataset and produces more biologically coherent outputs on real tissue, with iteration between decontamination and cell-cluster annotation further refining these outcomes.

We outline two limitations along with corresponding future work. First, DeSpotX is designed for single-cell-resolution ST platforms; spot-resolution platforms such as Visium or Slide-seq aggregate multiple cells per spot, and extending DeSpotX would require modeling cell-cluster mixtures. Second, the identifiability guarantee depends on the anchor genes within each cluster, which may be sparse in highly homogeneous tissues; future work could explore softer anchor formulations.

## 6 Acknowledgment

This work was supported by the National Institutes of Health NCI Cancer Systems Biology Consortium under award number U01CA264611.

## A Comparison of decontamination methods

Table 3 summarizes the key differences between DeSpotX and existing decontamination methods for scRNA-seq and ST data. The methods differ in the data type, the model architecture, the contamination estimator, the use of cluster annotations, and the constraint used to ensure identifiability. DeSpotX is the only method that introduces an explicit identifiability constraint through anchor genes, isolating spatial contamination from native expression in a cluster-aware framework.

**Table 3:**
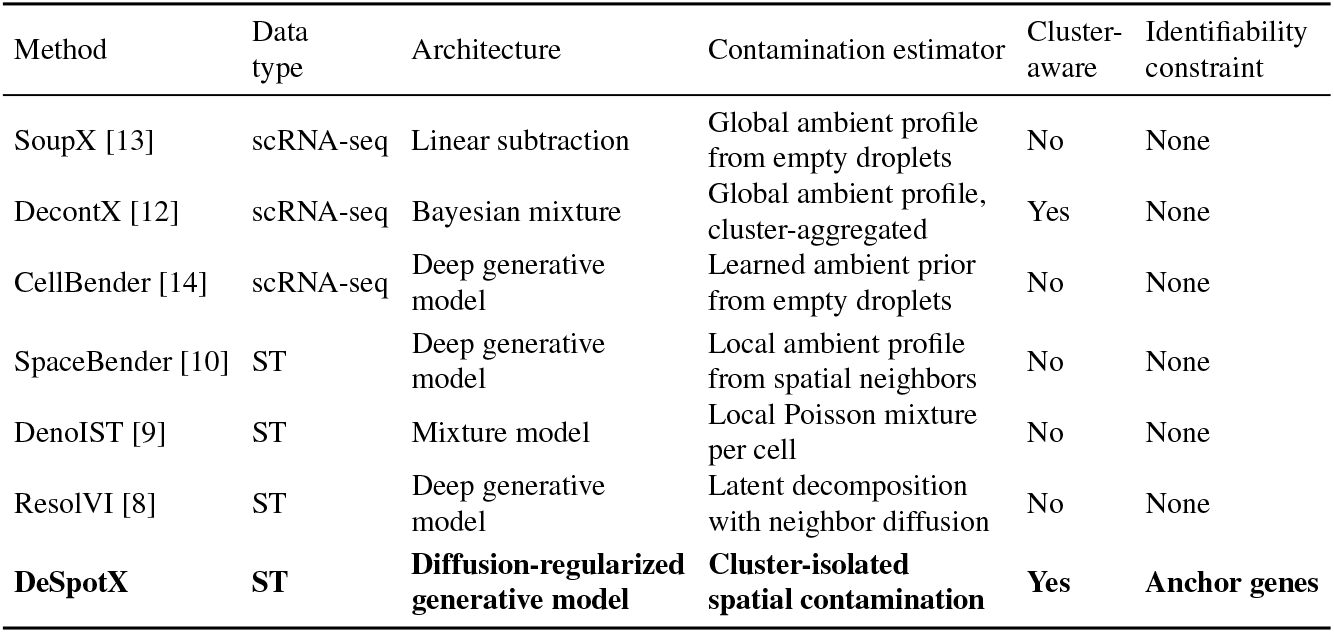
Comparison of decontamination methods for scRNA-seq and ST data.

| Method | Data type | Architecture | Contamination estimator | Cluster-aware | Identifiability constraint |
| --- | --- | --- | --- | --- | --- |
| SoupX [13] | scRNA-seq | Linear subtraction | Global ambient profile from empty droplets | No | None |
| DecontX [12] | scRNA-seq | Bayesian mixture | Global ambient profile, cluster-aggregated | Yes | None |
| CellBender [14] | scRNA-seq | Deep generative model | Learned ambient prior from empty droplets | No | None |
| SpaceBender [10] | ST | Deep generative model | Local ambient profile from spatial neighbors | No | None |
| DenoIST [9] | ST | Mixture model | Local Poisson mixture per cell | No | None |
| ResolVI [8] | ST | Deep generative model | Latent decomposition with neighbor diffusion | No | None |
| **DeSpotX** | **ST** | **Diffusion-regularized generative model** | **Cluster-isolated spatial contamination** | **Yes** | **Anchor genes** |

## B Proofs

### B.1 Proof of Lemma 1

We want to show two things: (i) the set 𝒮 (*m*) contains infinitely many triples, and (ii) the likelihood in Eq. (1) is the same at all of them.

Fix *m* ∈ Δ^*G−*1^. For any *ε* ∈ [0, 1) and any *χ* ∈ Δ^*G−*1^ satisfying *εχ*_*g*_ ≤ *m*_*g*_ for all *g*, define

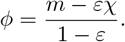

We check that the triple (*ε, ϕ, χ*) lies in 𝒮 (*m*), i.e., that it satisfies the conditions in the Lemma’s definition. First, *ϕ* is a valid probability vector; each coordinate is non-negative because 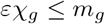, and the coordinates sum to one,

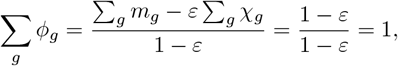

using Σ_*g*_ *m*_*g*_ = Σ_*g*_ *χ*_*g*_ = 1.

Second, *ϕ* satisfies the mixture equation: substituting the definition of *ϕ* gives

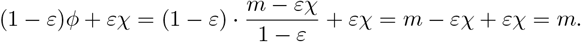

Hence (*ε, ϕ, χ*) ∈ 𝒮 (*m*).

Since *ε* and *χ* can be chosen from a continuum of admissible values, and each choice produces a distinct triple, 𝒮 (*m*) is a continuum, proving (i). For (ii), the likelihood in Eq. (1) depends on (*ε, ϕ, χ*) only through the mixture *m*, so it is constant across 𝒮 (*m*).

### B.2 Error under within-cluster contamination

DeSpotX’s cross-cluster mask (Eq. (3)) excludes same-cluster neighbors when estimating contamination, leaving a within-cluster residual in 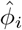. We show this residual contributes a native-profile error bounded by the contamination level times the within-cluster heterogeneity, while 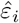 recovers the cross-cluster contamination rate.

Decompose the true contamination profile at cell *i* as

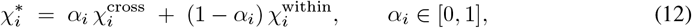

where *α*_*i*_ is the cross-cluster fraction. Substituting into Eq. (1) gives

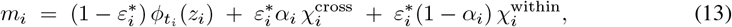

while DeSpotX fits

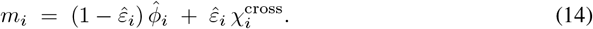

Let

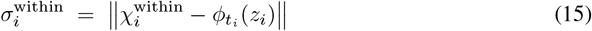

denote the within-cluster heterogeneity at cell *i*, the deviation between cell *i*’s native expression and the average expression of its same-cluster spatial neighbors.

#### Lemma 3

(Error from within-cluster contamination). *Under the conditions of Lemma 2, with* 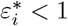, *define the scaling factor*

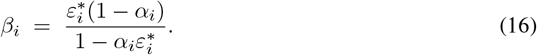

*The unique fit of Eq*. (14) *to the true mixture in Eq*. (13) *satisfies* 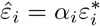 *and*

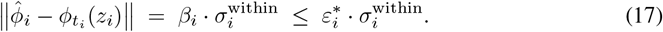

**Proof**. Equating Eqs. (13) and (14) gives

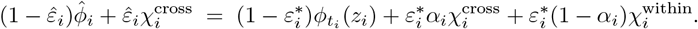

At any anchor position 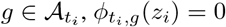 by definition. Since 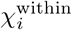 averages expression from cluster-*t*_*i*_ cells, which are zero at anchor positions, 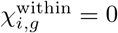 as well. The anchor constraint forces 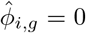, so the equation at *g* reduces to 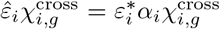. Since at least one anchor satisfies 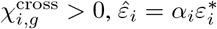. Substituting back,

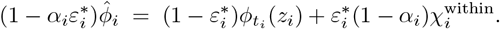

Dividing by 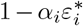 and subtracting 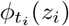 gives 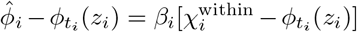, and Eq. (17) follows by taking norms.

#### Remarks

The error in Eq. (17) is the product of two small quantities. The scaling factor *β*_*i*_ is bounded by 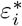(for example, *β*_*i*_ = 0.07 at 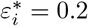 and *α*_*i*_ = 0.7). The within-cluster heterogeneity 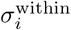 is small for well-defined clusters.

The recovered contamination fraction 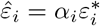 is the cross-cluster rate, which drives the downstream artifacts that motivate decontamination, such as cell-type misannotation, smeared marker maps, and spurious cell-cell communication. The within-cluster portion absorbed into 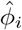 does not cause these artifacts, since the recipient cell expresses similar genes natively.

We defer validation of Lemma 3 on spike-in simulations to Appendix I.

### B.3 Proof of Lemma 2

We show two things: (i) *ε*_*i*_ is identifiable from observations at anchor positions, and (ii) 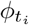 is identifiable at all non-anchor positions. Both follow under the preconditions of the Lemma: *χ*_*i*_ given as input, constructed without using *x*_*i*_, and at least one anchor with *χ*_*i,g*_ *>* 0.

Recall that 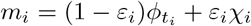 denotes the mixture mean satisfying 𝔼[*x*_*i*_*/d*_*i*_] = *m*_*i*_. Because *χ*_*i*_ is not a function of *x*_*i*_, the identification of *ε*_*i*_ and 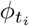 from *x*_*i*_ is not circular.

For (i), consider any anchor position 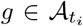. By the anchor assumption 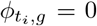, the mixture equation reduces to

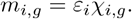

Since by assumption there exists 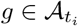 with *χ*_*i,g*_ *>* 0, solving gives

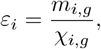

which is a function of observable quantities and the (independently obtained) *χ*_*i*_.

For (ii), fix *ε*_*i*_ and *χ*_*i*_ as above. For any non-anchor position 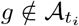, the mixture equation gives

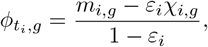

which expresses 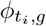 in terms of observables and parameters identified in (i).

Hence both *ε*_*i*_ and 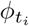 are identifiable.

## C Model details

### C.1 Spatial graph encoder: implementation details

#### Node features

For each node *j* in the star graph 𝒢_*i*_, the input feature is

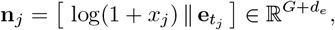

where log(1 +·) is applied elementwise to the count vector, 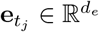 is a learned cluster embedding, and ∥ denotes concatenation. The cluster embedding is initialized randomly and trained jointly with the rest of the model. A linear projection 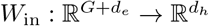 to a hidden representation 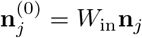 used as input to the GAT layer.

#### Edge features

The edge from leaf *j* to center *i* carries the kernel weight *w*_*ij*_ = exp(−∥*s*_*j*_ − *s*_*i*_ */ρ*_*i*_) ∈ (0, 1] as a single scalar feature, with *ρ*_*i*_ set to the median Euclidean distance from cell *i* to its *K* nearest neighbors. The exponential form approximates the spatial decay of free ambient RNA, and the per-cell adaptive bandwidth *ρ*_*i*_ accommodates variation in cell density across the dataset.

#### GATv2 attention

GATv2 [18] computes attention coefficients of the form

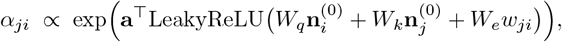

where *W*_*q*_, *W*_*k*_ are query and key projections, *W*_*e*_ embeds the scalar edge weight *w*_*ji*_, and **a** is a learned scoring vector. We use *H* attention heads with averaging across heads and per-head output dimension *d*_*h*_, so the output dimension is *d*_*h*_. For batched training, *B* star graphs are concatenated into a single disjoint graph and the center nodes are recovered by index after the forward pass.

#### Readouts from *h*_*i*_

Two linear heads applied to the center’s spatial-context embedding *h*_*i*_ produce the latent state and the contamination-fraction estimate in Eq. (2):

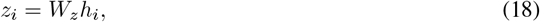

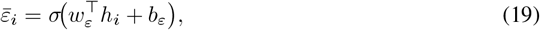

where *W*_*z*_, *w*_*ε*_, and *b*_*ε*_ are learned parameters. We initialize *b*_*ε*_ = logit(0.20).

#### Hyperparameters

Default values used in our experiments: GAT hidden dimension *d*_*h*_ = 256, *H* = 4 attention heads, cluster embedding dimension *d*_*e*_ = 32, *K* = 12 spatial neighbors, and dropout 0.1.

### C.2 Latent diffusion prior: implementation details

#### Forward diffusion

Following the standard DDPM formulation [19], we define a discrete-time forward diffusion on *z* with *T* steps and a fixed variance schedule 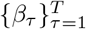. The marginal at step *τ* is

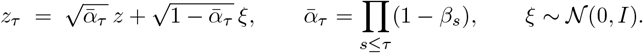

#### Score network

The score network *s*_*ω*_ is a feedforward MLP whose inputs are the noised latent *z*_*i,τ*_, a sinusoidal embedding of the diffusion step *τ*, the spatial-context embedding *h*_*i*_, and a learned cluster embedding indexed by *t*_*i*_. Its output is a prediction of the noise *ξ* added at step *τ* .

#### Stop-gradients

The stop-gradient operator on *h*_*i*_ and *z*_*i,τ*_ in Eq. (5) blocks diffusion-loss gradients from reaching the encoder, so the encoder is shaped only by the reconstruction loss. This prevents a degenerate solution in which the encoder collapses *z*_*i*_ into a distribution that *s*_*ω*_ trivially denoises, leaving the prior uninformative. It also ensures that the prior tracks the encoder’s reconstruction-driven distribution, so refinement at inference projects 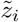 toward the same distribution the decoder was trained on.

#### Inference-time refinement

At inference, we noise the encoder output *z*_*i*_ to diffusion step *τ*_start_ = ⌊*λ*_refine_ · *T* ⌋ and run *K*_refine_ DDIM reverse steps [20] to obtain a refined latent 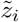, which is decoded to yield 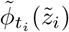.

#### Hyperparameters

The score network is a 3-layer MLP with hidden dimension 256 and SiLU activation. The forward diffusion uses *T* = 100 steps with linear schedule *β*_*τ*_ from 10^*−*4^ to 0.02. At inference, we use noise fraction *λ*_refine_ = 0.2 and *K*_refine_ = 10 DDIM steps.

### C.3 Cluster-conditioned decoder: implementation details

#### Architecture

The function *f*_*θ*_ in Eq. (6) is a two-layer MLP with hidden dimension *d*_dec_ and Softplus activations:

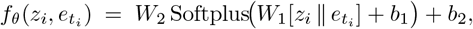

where [· ∥ ·] denotes concatenation, 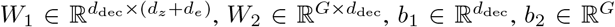, and 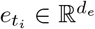 is a learned cluster embedding. We initialize *W*_2_ with Xavier-uniform values scaled by 0.1 and *b*_2_ = 0.

#### Cluster-specific logit shift

The vector 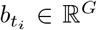 in Eq. (6) is initialized to 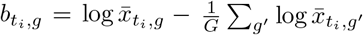, where 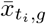 is the mean count of gene *g* over cluster-*t*_*i*_ training cells. After initialization, 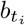is fixed; cluster-specific learnable signal is carried by 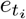 and *f*_*θ*_.

#### Hyperparameters

Default values used in our experiments: *d*_dec_ = 256, *d*_*e*_ = 32.

### C.4 Anchor mask construction: implementation details

#### Per-cluster expression statistics

Let *r*_*t,g*_ denote the fraction of cluster-*t* cells with nonzero counts for gene *g* in the training data. We construct the binary anchor mask

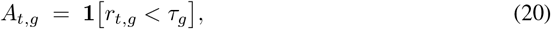

with adaptive per-gene threshold

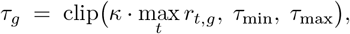

which scales with each gene’s maximum cluster-level expression rate. Default value used in our experiments: *κ* = 0.3.

### C.5 Training: implementation details

#### Loss weights

Default values used in our experiments: *λ*_*A*_ = 50, *λ*_*D*_ = 1, *λ*_*E*_ = 1. Although *λ*_*A*_ appears large relative to *λ*_*D*_ and *λ*_*E*_, the anchor penalty vanishes once 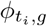 is driven to zero at anchor positions, so *λ*_*A*_ only governs the convergence speed of this collapse.

#### Regularizer on 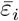

The Beta-density form in Eq. (10) provides a smooth, bounded penalty whose minimum lies at *µ*_*ε*_. We use fixed concentration *v* = 20 and treat *µ*_*ε*_ as a single global scalar, parameterized by its logit and initialized at logit(0.20).

#### Anchor warm-up

The anchor weight *λ*_*A*_ is linearly increased from 0 to 50 over the first *E*_warm_ = 3 epochs and held at 50 thereafter.

#### Optimizer

All parameters, including the encoder, decoder, score network, gene-wise dispersion *θ*, and *µ*_*ε*_, are optimized with Adam (*β*_1_ = 0.9, *β*_2_ = 0.999). We use learning rate 10^*−*3^, batch size 128, and gradient clipping at norm 10. Training runs for 10 epochs.

#### Compute

All experiments were run on a single NVIDIA Tesla T4 GPU (16 GB), with training taking 5–20 minutes per dataset.

## D Spike-in simulation details

### Ambient profile c onstruction

For each cell *i* at spatial coordinate **s**_*i*_, with native counts **n**_*i*_ ∈ ℕ^*G*^, library size *d*_*i*_ = Σ_*g*_ *n*_*i,g*_, and cell-cluster label *t*_*i*_, we construct a per-cell ambient profile

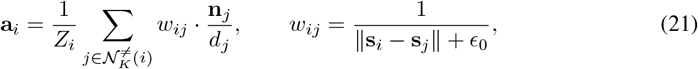

where 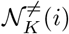 is the set of *K* = 12 Euclidean nearest neighbors of cell *i* restricted to cells with *t*_*j*_ ≠ *t*_*i*_, falling back to the unrestricted neighbor set when no other-cluster neighbor exists; *ϵ*_0_ = 10^*−*6^ is a numerical stabilizer; and 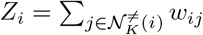 normalizes **a**_*i*_ to the simplex.

### Per-cell contamination rates

We draw per-cell contamination rates from a Beta distribution centered at the noise level 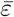,

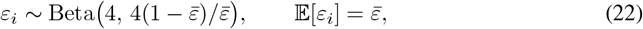

with shape parameter fixed at 4 to give moderate per-cell variance.

### Spike-in count sampling

Synthetic contamination counts are sampled independently per gene as

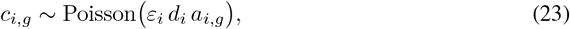

and the spiked counts delivered to each method are *s*_*i,g*_ = *n*_*i,g*_ + *c*_*i,g*_.

## E Ablation studies

We perform ablation studies on `Xenium_Breast` and `MERFISH_Brain` to assess the contribution of each architectural component. We compare the full DeSpotX model against four ablations, each of which removes one component while keeping the rest of the architecture intact.

### Ablation conditions

Each ablation removes a single component from the full model.

- *w/o spatial*: spatial information is removed. The GAT layer is replaced by an MLP applied to each cell’s own counts, and *χ*_*i*_ is replaced by a single dataset-level ambient profile shared across cells.
- *w/o cluster-mask*: cluster conditioning is removed. The cross-cluster mask in *χ*_*i*_ (Eq. 3) is dropped so all neighbors contribute regardless of cluster identity, and the cluster-specific decoder bias 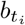 is no longer warm-started from per-cluster expression statistics.
- *w/o diffusion*: the latent diffusion prior is disabled. The diffusion loss term is set to zero during training (setting *λ*_*D*_ = 0), and the DDIM refinement step is skipped at inference, so the encoder latent *z*_*i*_ is decoded directly.
- *w/o identifiability*: identifiability constraints are removed. This includes the anchor penalty (setting *λ*_*A*_ = 0), the Beta prior on 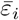 (setting *λ*_*E*_ = 0), and the architectural priors that contribute to identifiability. With these constraints removed, the negative-binomial likelihood and the simplex constraint on *ϕ* remain, corresponding to the unconstrained setting of Lemma 1.

### Results

Figure 5 shows that the full model achieves the best mean performance on every metric and dataset, indicating that each architectural component contributes to DeSpotX’s overall performance. The largest effect comes from removing the identifiability constraints. On `Xenium_Breast`, PCE rises from a median of 16 to 60 pp and GCE rises from 10 to above 70 pp, with both metrics also showing greatly increased variance across noise levels. AUROC drops substantially and becomes highly variable. On `MERFISH_Brain`, the same ablation raises PCE from a median of 10 to 20 pp and GCE from 5 to 17 pp, with AUROC dropping to a median of *<*0.90. These results are consistent with Lemma 1, which states that without external constraints, the (*ϕ, ε, χ*) decomposition is not uniquely determined.

**Figure 5:**
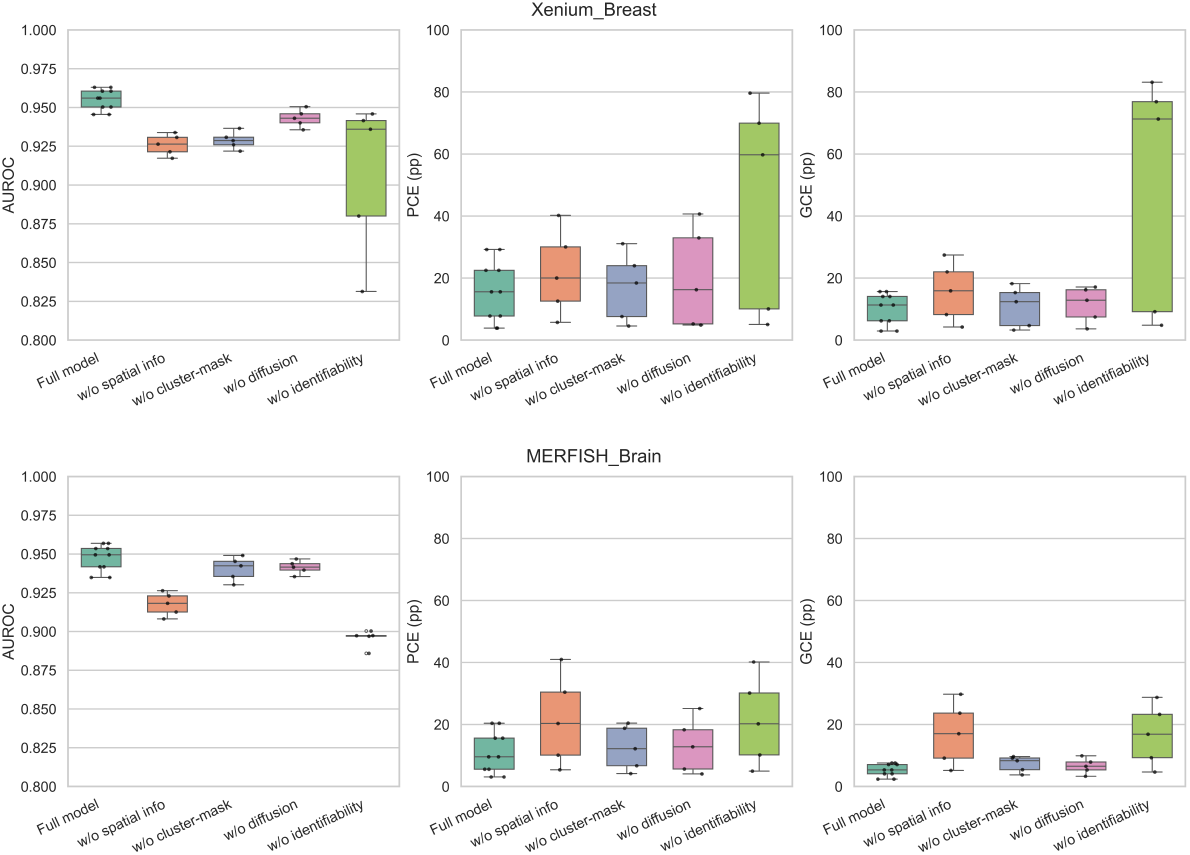
Ablation studies on `Xenium_Breast` (top row) and `MERFISH_Brain` (bottom row), reporting AUROC, PCE, and GCE across the five non-zero noise levels. Each box summarizes the distribution of values across noise levels. Higher AUROC is better; PCE and GCE are reported in percentage points and lower is better.

Removing the spatial information produces consistent degradation across all three metrics on both datasets, confirming that the spatial graph encoder and the local contamination estimator are central to DeSpotX’s accuracy. Removing the cluster-mask and the diffusion prior also degrades all three metrics on both datasets, with smaller effects than the spatial ablation. The cluster-mask supplies cell-cluster information that helps separate native from contaminating signal, whereas the diffusion prior reduces distortions introduced by the contamination estimator; the results confirm that both components contribute to DeSpotX’s performance.

## F Effect of the diffusion prior on low-but-real expression

We test whether DeSpotX’s diffusion prior helps preserve low-but-real expression of established marker genes. We use the spike-in benchmark data at 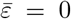 (no injected contamination) on `Xenium_Breast` and `MERFISH_Brain` datasets, and train DeSpotX with three random seeds, with and without the diffusion prior. All other hyperparameters are held fixed across paired runs so that the only difference is the diffusion contribution.

We focus on established, biologically validated markers expressed at low levels in their native cell-clusters, where any removal by a decontamination model represents over-correction of real signal. Specifically, we select markers with intrinsic mean count *<* 3 in their native cell-clusters. For `MERFISH_Brain`, this yields 12 brain markers including neurotransmitter enzymes (Th, Chat, Nos1), neuropeptides (Crh, Vip, Gal, Pdyn, Tac2), dopamine receptors (Drd1, Drd2), an MGE transcription factor (Sox6), and calretinin (Calb2), with native means in the range 0.4 to 2.7. For `Xenium_Breast`, this yields 6 breast immune and proliferation markers, including MKI67 in proliferating tumor cells, ITGAX and C1QA in macrophages and dendritic cells, and CD27, GZMB, CTLA4 in T cells, with native means in the range 0.1 to 2.1.

For each (cell, marker) pair where the cell is native to that marker and the input count is positive, we compute retention 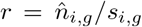. We then report the per-marker change in retention Δ*r* = *r*_with_ − *r*_without_ between runs with and without the diffusion prior, in percentage points. Positive values indicate that the diffusion prior helps preserve more native signal at low expression levels.

Figure 6 shows the per-marker Δ*r* on both datasets. On `MERFISH_Brain`, every one of the 12 markers shows positive Δ*r*, with values from +0.33 to +1.55 percentage points. On `Xenium_Breast`, 4 of 6 markers show positive Δ*r* and 2 are neutral, with the largest gains on MKI67 (+0.34) and ITGAX (+0.23). These results indicate that the diffusion prior helps preserve low-expression marker-gene signal.

**Figure 6:**
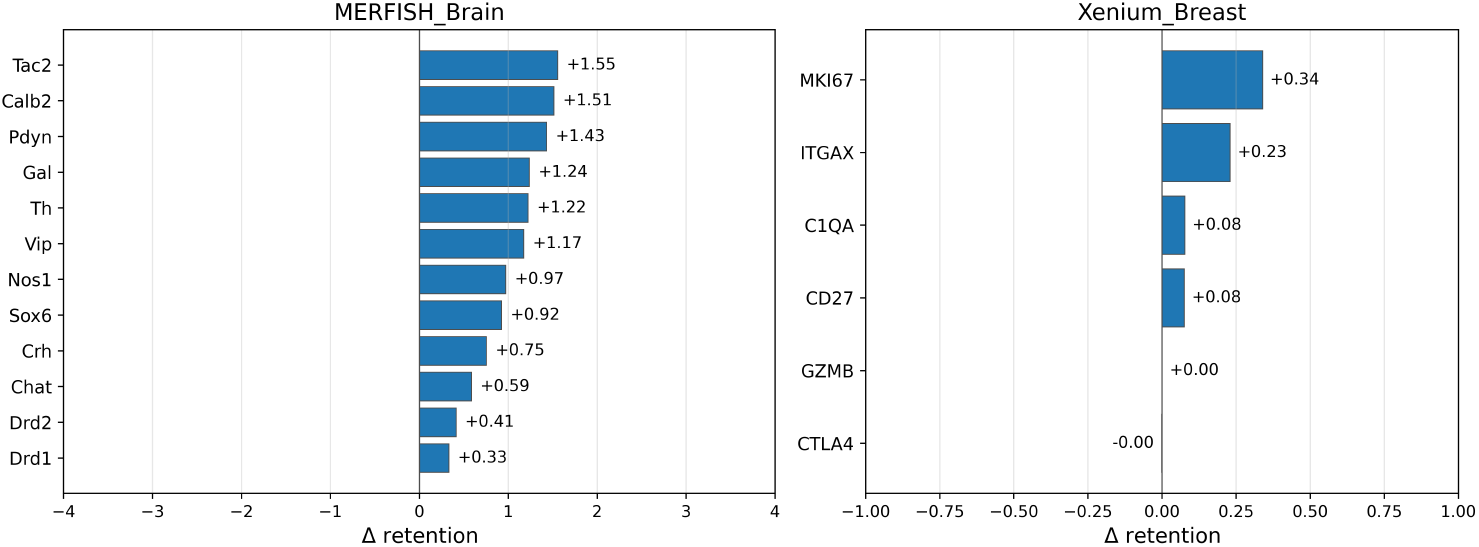
Effect of the diffusion prior on retention of low-but-real expression. Per-marker change in retention Δ*r* between DeSpotX runs with and without the diffusion prior. Positive values indicate that the diffusion prior helps preserve more native signal.

## G Robustness studies

We assess DeSpotX’s robustness to perturbations of five inputs: the adaptive-anchor threshold *κ*, random perturbation of the anchor mask, noise in the cell-cluster labels, the number of spatial neighbors *K*, and the cell-cluster annotation method. For each factor, we run the spike-in benchmark on `Xenium_Breast` and `MERFISH_Brain` with the perturbed input while keeping the rest of the configuration at its default.

### Robustness conditions

Each condition perturbs a single input from the default configuration.

- *Adaptive-anchor threshold*: *κ* is varied across {0.1, 0.2, 0.3, 0.4, 0.5} . Smaller *κ* produces a more permissive threshold and anchors more genes per cluster.
- *Anchor mask perturbation*: a uniform fraction {5%, 10%, 20%} of entries in the default *κ* = 0.3 anchor mask is randomly flipped (both 0 → 1 and 1 → 0).
- *Cell-cluster label perturbation*: the published cell-cluster annotation is perturbed by reassigning a random fraction {5%, 10%, 20%, 30%} of cells to a different cluster, sampled uniformly. Both the cross-cluster mask in *χ*_*i*_ (Eq. 3) and the anchor mask (Eq. 20) are derived from the cluster annotation.
- *Number of spatial neighbors*: *K* is varied across {8, 12, 24} . *K* sets the number of spatial neighbors used by the GAT graph and the contamination estimator *χ*_*i*_ (Eq. 3).
- *Cell-cluster annotation method*: the published cell-cluster annotation is replaced by Leiden clustering at three target cluster counts *N* ∈ { 15, 25, 30} . The Leiden resolution is binary-searched per dataset to hit each target.

#### Results

DeSpotX is robust to perturbations of the anchor mask. Across *κ* ∈ {0.1, 0.2, 0.3, 0.4, 0.5}, AUROC moves by at most 0.011 on either dataset, with PCE and GCE remaining close to their default values (Figure 7). When up to 20% of mask entries are flipped, no metric degrades on either dataset (Figure 8). DeSpotX does not require complete accuracy of the anchor mask, as identification can be supported by a sufficient subset of correct anchors within each cluster. This makes DeSpotX tolerant of moderate inaccuracies in the threshold *κ* and in the anchor mask.

**Figure 7:**
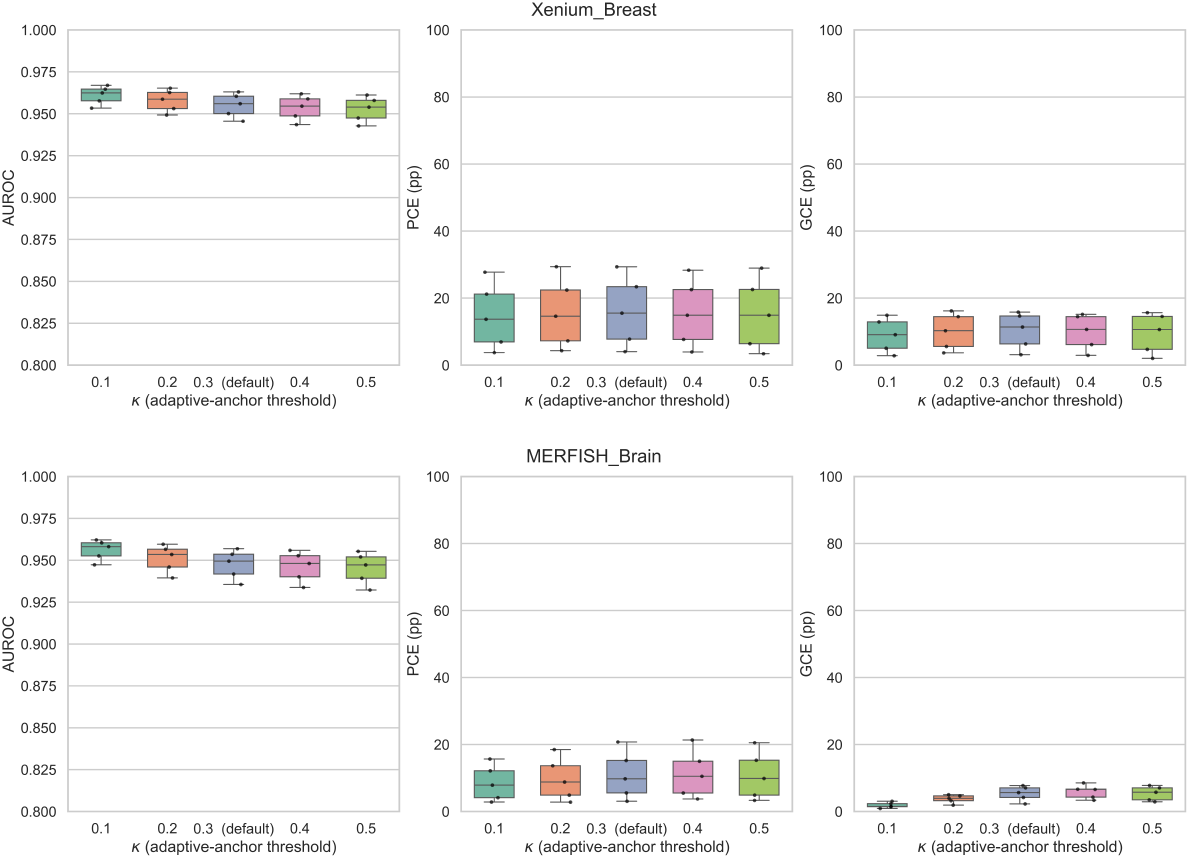
Robustness to the adaptive-anchor threshold *κ*. Each box summarizes the metric distribution across the five non-zero noise levels. PCE and GCE are reported in percentage points.

**Figure 8:**
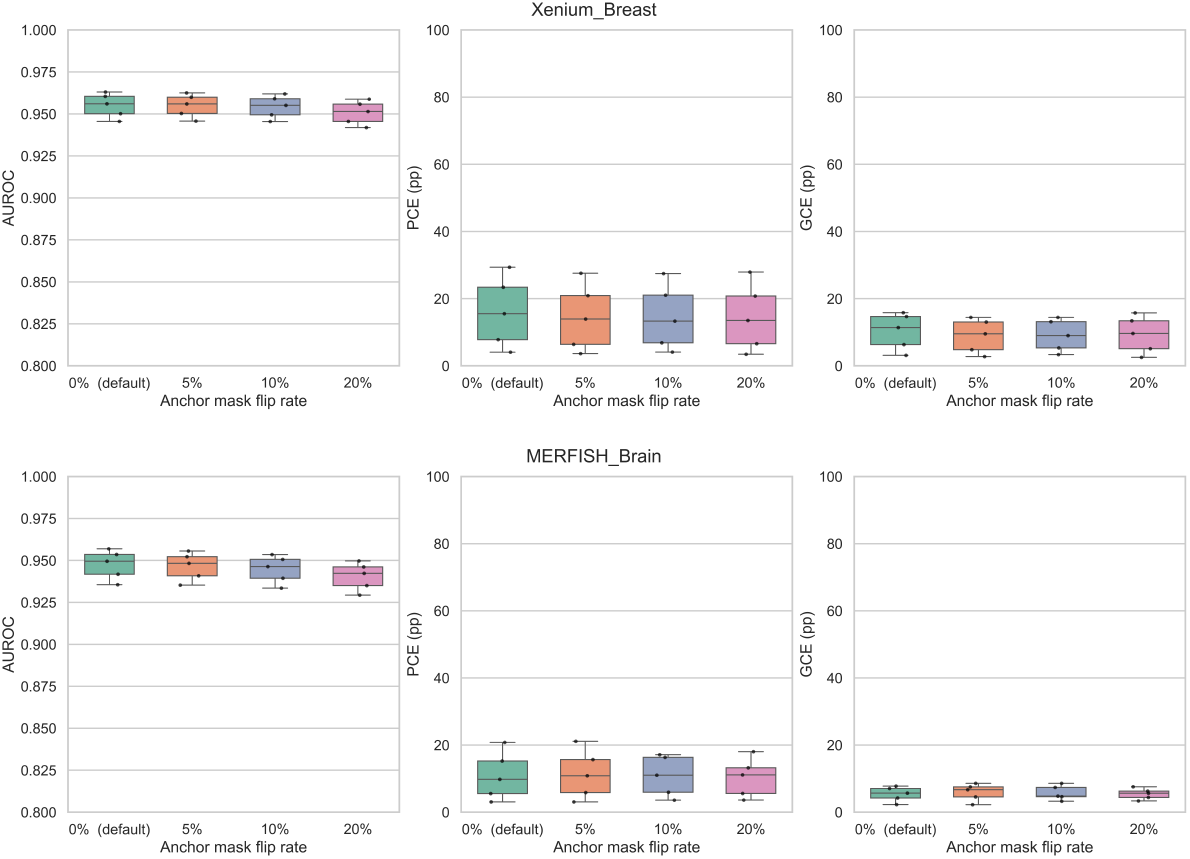
Robustness to random perturbation of the anchor mask. Each box summarizes the metric distribution across the five non-zero noise levels. PCE and GCE are reported in percentage points.

Figure 9 shows that all three metrics remain close to the default values as the cell-cluster label perturbation rate increases from 0% to 30%. AUROC drops by less than 0.005 on both datasets, indicating that DeSpotX is not sensitive to label noise. Figure 10 shows that all three metrics remain close to the default values across *K* ∈ {8, 12, 24} on both datasets, indicating that DeSpotX is robust to the choice of *K* in this range.

**Figure 9:**
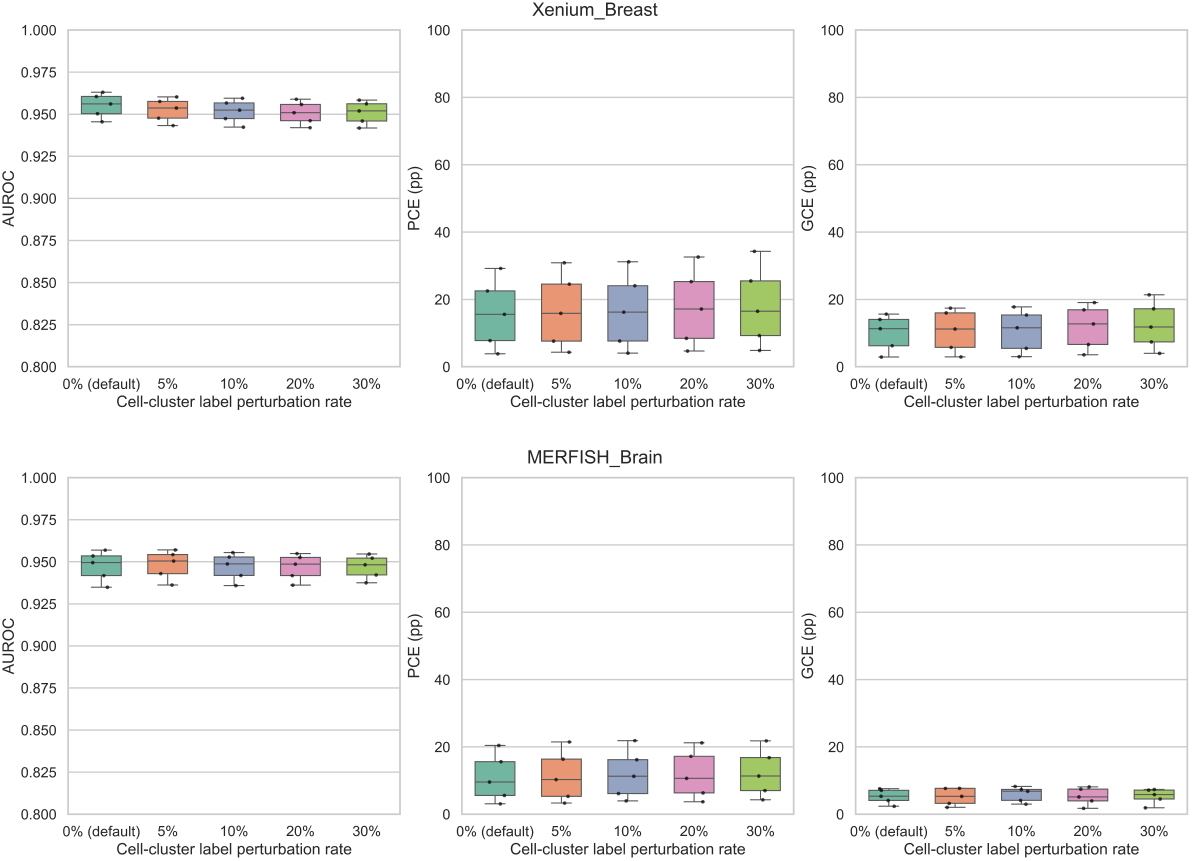
Robustness to cell-cluster label perturbation. Each box summarizes the metric distribution across the five non-zero noise levels. PCE and GCE are reported in percentage points.

**Figure 10:**
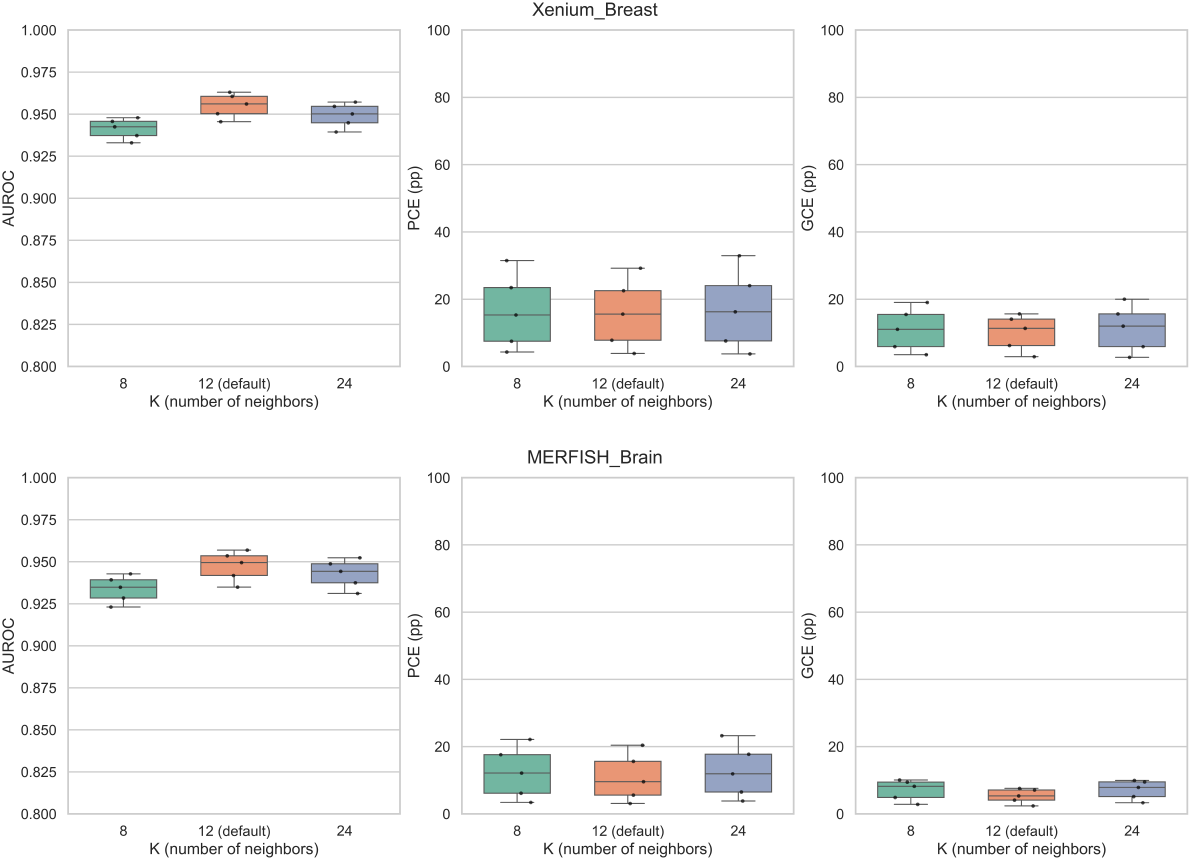
Robustness to the number of spatial neighbors *K*. Each box summarizes the metric distribution across the five non-zero noise levels. PCE and GCE are reported in percentage points.

When the published cell-cluster annotation is replaced by Leiden clustering, AUROC drops by less than 0.03 on both datasets, while PCE and GCE remain close to the default values (Figure 11). The performance is consistent across the three Leiden cluster counts *N* ∈ {15, 25, 30}, indicating that DeSpotX is not sensitive to the specific cluster count chosen by Leiden.

**Figure 11:**
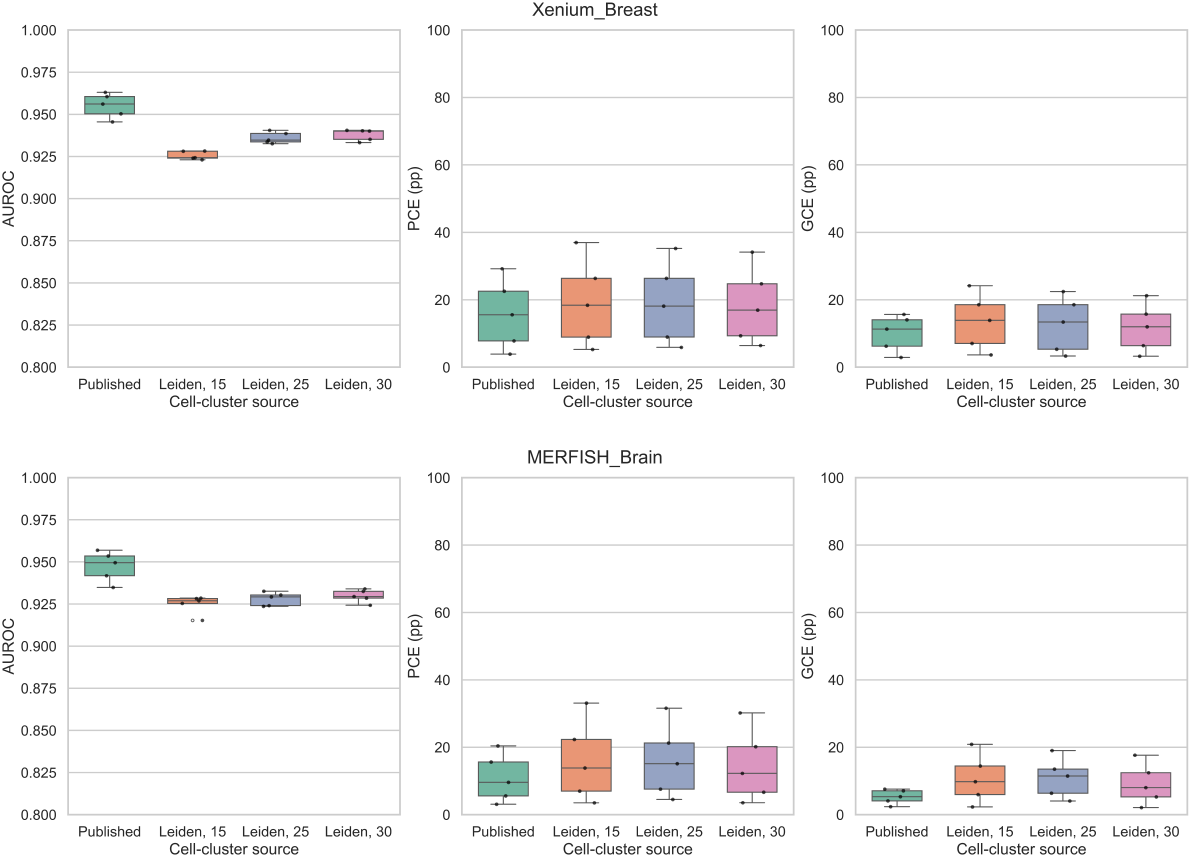
Robustness to the cell-cluster annotation method. Each box summarizes the metric distribution across the five non-zero noise levels. PCE and GCE are reported in percentage points.

## H Effect of contamination level on performance

We examine how DeSpotX’s performance varies with the noise level *ε* on each of the five spike-in benchmark datasets.

### AUROC of DeSpotX across noise levels

Figure 12 shows DeSpotX’s AUROC at each noise level on each dataset. Across all five datasets, AUROC varies by at most 0.025 between 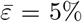 and 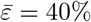. On `Xenium_Breast` and `MERFISH_Brain`, AUROC increases slightly with 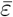, since stronger contamination produces clearer signals that are easier to identify against the native expression background. DeSpotX therefore performs consistently across contamination levels.

**Figure 12:**
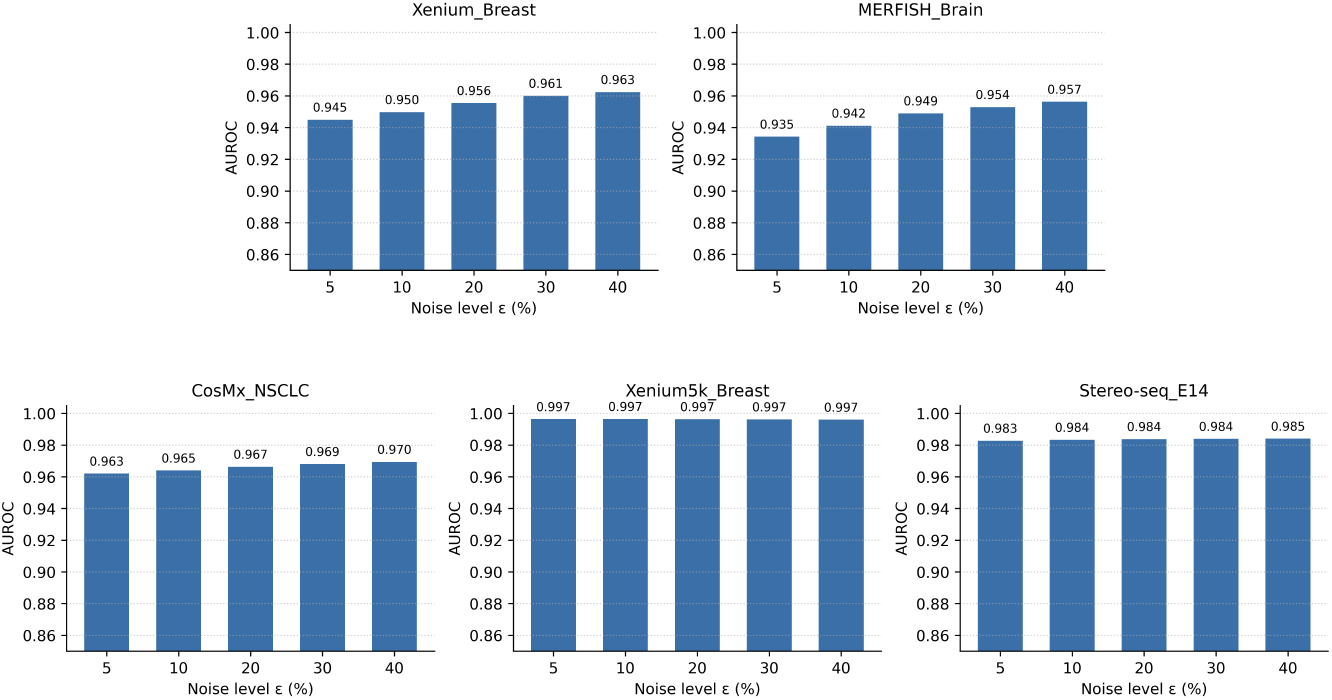
DeSpotX’s AUROC on each of the five spike-in benchmark datasets at five contamination levels *ε* ∈ {5, 10, 20, 30, 40}%.

### Per-cell calibration advantage of DeSpotX across noise levels

We compare DeSpotX against DecontX and ResolVI as representative baselines for scRNA-seq and ST methods, respectively. Figure 13 reports the per-cell calibration error gap ΔPCE = PCE_baseline_ − PCE_DeSpotX_ between DeSpotX and these two baselines, in percentage points. Positive values indicate that DeSpotX achieves lower calibration error than the baseline. On every dataset and at every 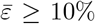, both gaps are positive, and the gaps grow with 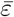. At 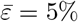, the gaps are within 3 pp; at 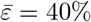, the gaps reach 11–13 pp on `CosMx_NSCLC` and `MERFISH_Brain` and 22–27 pp on `Xenium5k_Breast`. The increasing gap shows that DeSpotX’s per-cell contamination estimates remain accurate as the contamination signal grows, while baseline calibration error scales with the contamination level.

**Figure 13:**
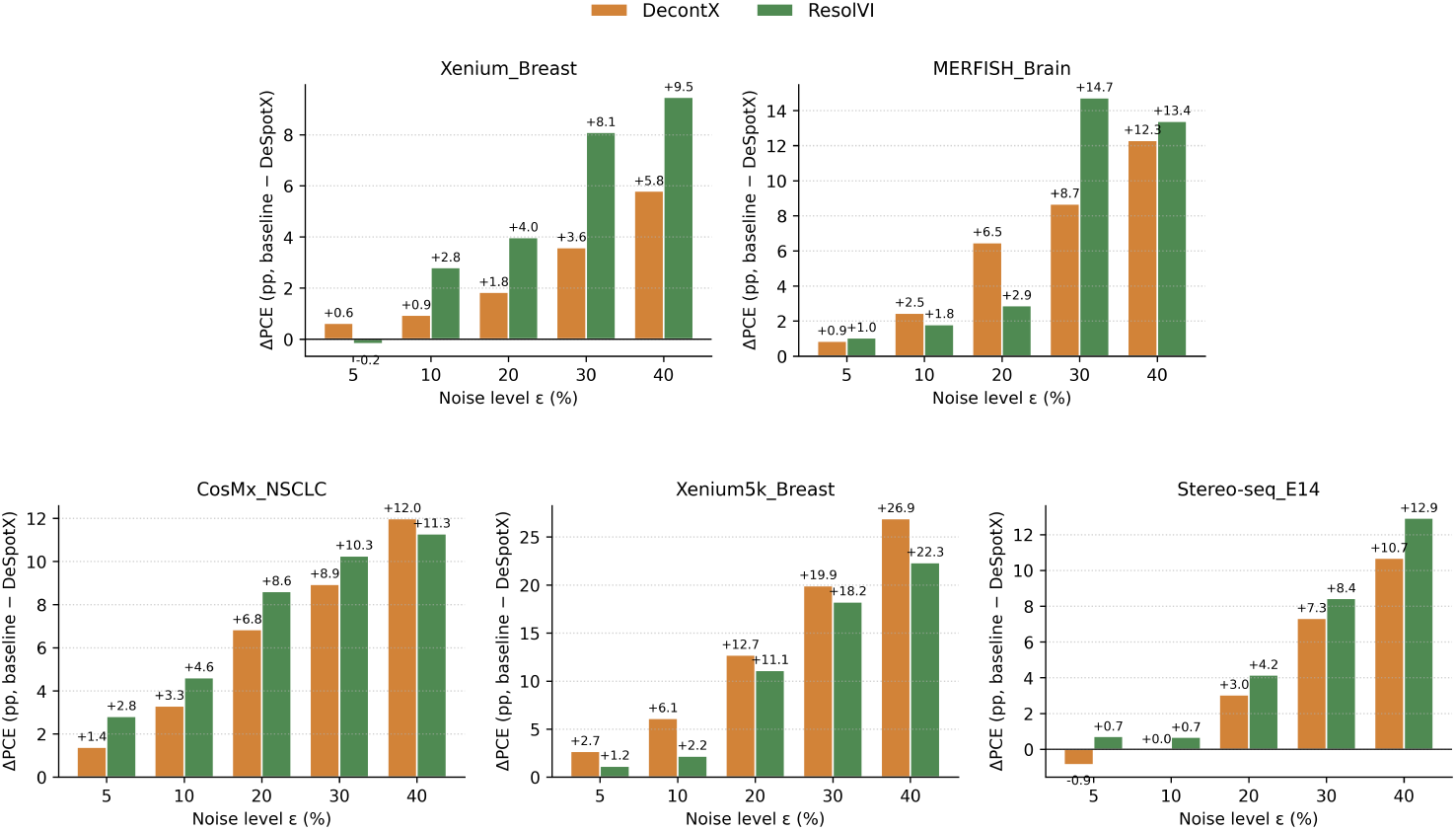
Per-cell calibration error gap ΔPCE = PCE_baseline_ − PCE_DeSpotX_ between DeSpotX and the representative baselines (DecontX, ResolVI), in percentage points, at five contamination levels 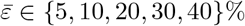.

## I Validation of the within-cluster bound

We test Lemma 3 (Appendix B.2) on a spike-in simulation in which each cell receives contamination from a mixture of its cross-cluster and same-cluster spatial neighbors, weighted *α* and 1 − *α* respectively, with per-cell rate sampled from a Beta distribution centered at 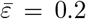. We vary *α* ∈ {0.3, 0.5, 0.7, 0.9, 1.0} on `Xenium_Breast` and `MERFISH_Brain`.

AUROC on cross-cluster contamination labels stays above 0.93 across all *α* on both datasets (Figure 14a), confirming 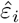 tracks the cross-cluster rate. The per-cell native-profile error grows linearly with the scaling factor *β*_*i*_, with slopes 0.015 on `Xenium_Breast` and 0.010 on `MERFISH_Brain` (Figure 14b), well below the empirical ⟨*σ*^within^⟩ of 0.057 and 0.063 measured across the five datasets (Figure 14c). The intercept (≈0.022) reflects *α*-independent error sources unrelated to within-cluster contamination, including DeSpotX’s removal of natural platform contamination from the unspiked reference.

**Figure 14:**
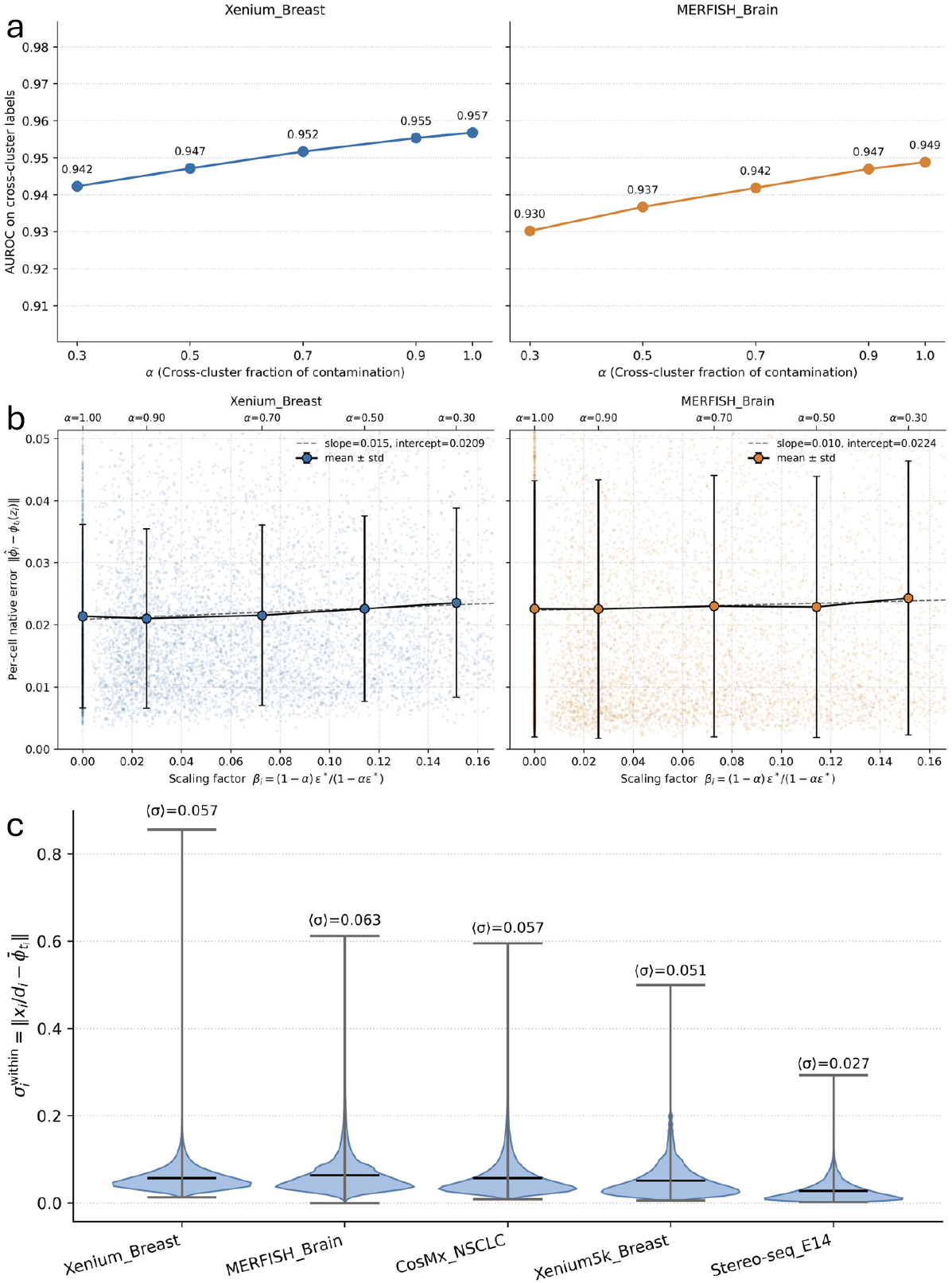
DeSpotX is robust to within-cluster contamination across spike-in mixtures. (a) AUROC for distinguishing cross-cluster contamination from native counts, across *α* ∈ {0.3, 0.5, 0.7, 0.9, 1.0}. (b) Per-cell native-profile error 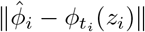 versus the scaling factor *β*_*i*_ = (1 − *α*)*ε*^*\**^*/*(1 − *αε*^*\**^). Markers show mean ± std per *α*; dashed line is a linear fit. (c) Distribution of 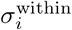 across cells in all five datasets at zero injected contamination, with mean values annotated.

## J Runtime comparison

We compare end-to-end runtime across decontamination methods on `Xenium5k_Breast` (577,258 cells, 5,101 genes) and `Stereo-seq_E14` (92,928 cells, 18,582 genes), the two largest datasets used in this work (Figure 15). SoupX and DecontX run in 3 to 10 minutes. Among the deep learning-based methods, DeSpotX is the fastest at 16 to 21 minutes. ResolVI and SpaceBender take 6 to 53 hours. All methods were run on a single NVIDIA Tesla T4 GPU (16 GB).

**Figure 15:**
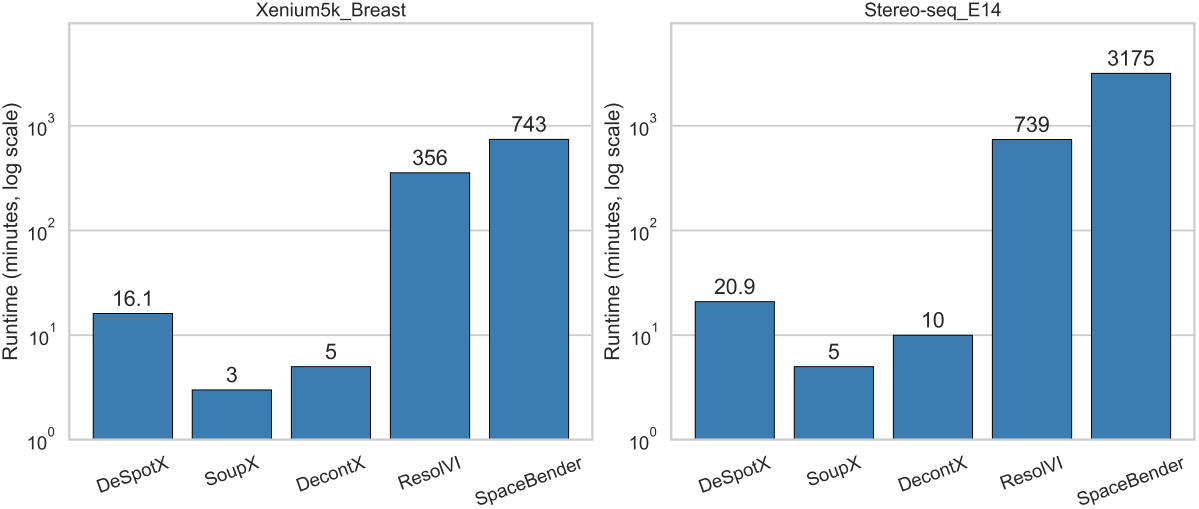
Runtime comparison across decontamination methods on `Xenium5k_Breast` and `Stereo-seq_E14`.

## K Marker-gene specificity of baseline methods

We extend the marker-gene comparison in Figure 2b to the four baseline methods (Figure 16). SoupX produces dotplots that closely resemble the raw counts, with off-target expression largely unchanged. DecontX and SpaceBender reduce off-target expression to varying degrees across datasets, but residual signal in non-canonical clusters remains visible, particularly on the larger panels (Xenium5k_Breast, CosMx_NSCLC). ResolVI shows little difference between canonical and non-canonical clusters. All four baselines yield lower marker-gene specificity than DeSpotX.

**Figure 16:**
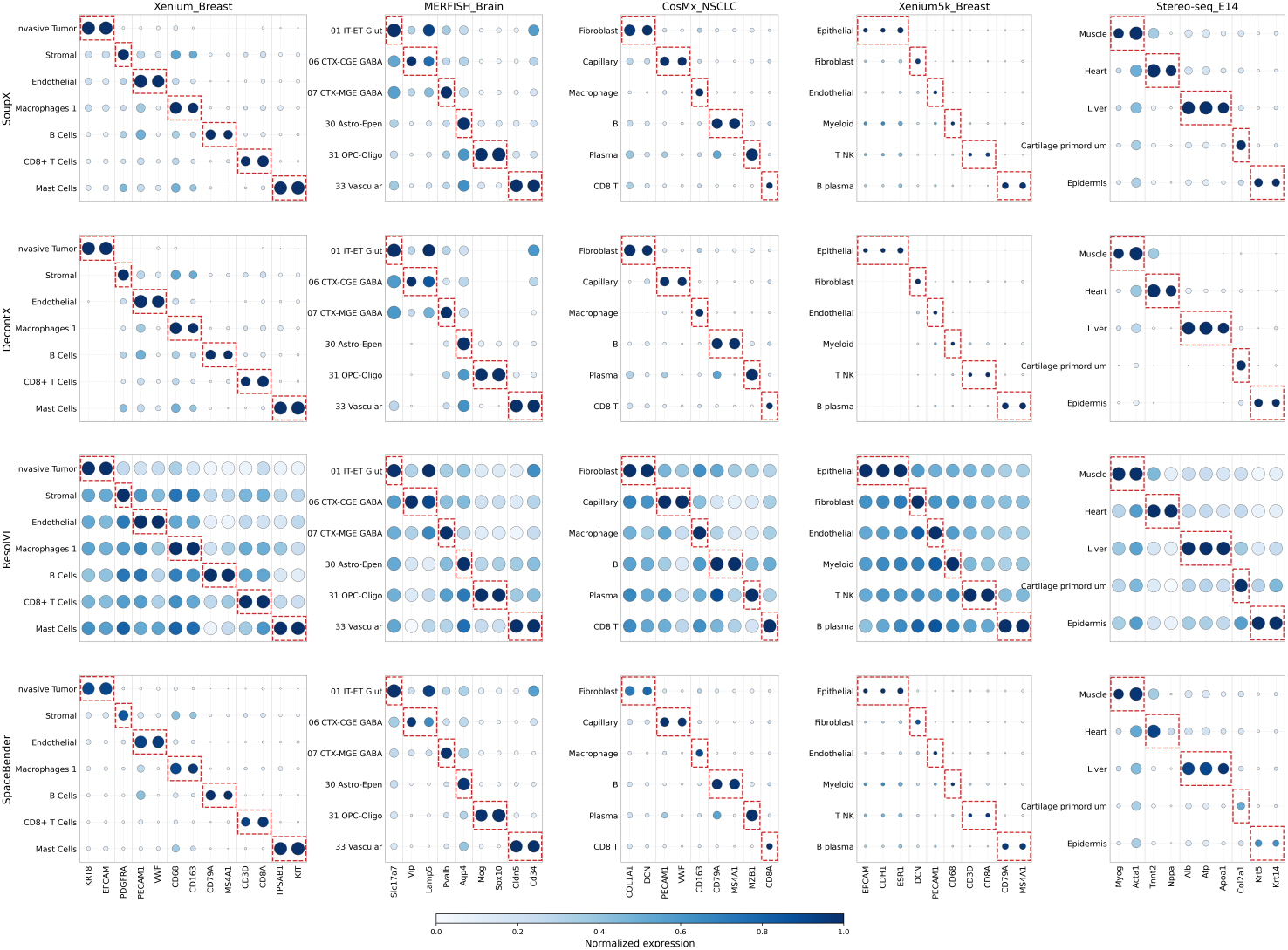
Marker-gene dotplots for the four baseline methods across the five datasets. Red dashed boxes mark canonical marker-cluster pairs.

## L Spatial localization of removed contamination

To examine where DeSpotX removes contamination, we map the per-cell amount removed for canonical markers of representative cell clusters across the five datasets (Figure 17). The removed counts are spatially concentrated near the canonical expressing cluster, consistent with the expected pattern of ambient contamination, in which a marker’s signal diffuses from cells that natively express it into nearby cells, and DeSpotX removes the diffused signal from the receiving cells. For example, VWF in `Xenium_Breast` is endothelial-specific and Mog in `MERFISH_Brain` is oligodendrocyte-specific; in both cases, DeSpotX’s removal extends beyond the canonical cluster into surrounding tissue, with the highest removal in regions immediately adjacent to the canonical cells. This pattern holds across the five datasets and supports the spatial-locality assumption underlying DeSpotX’s contamination model.

**Figure 17:**
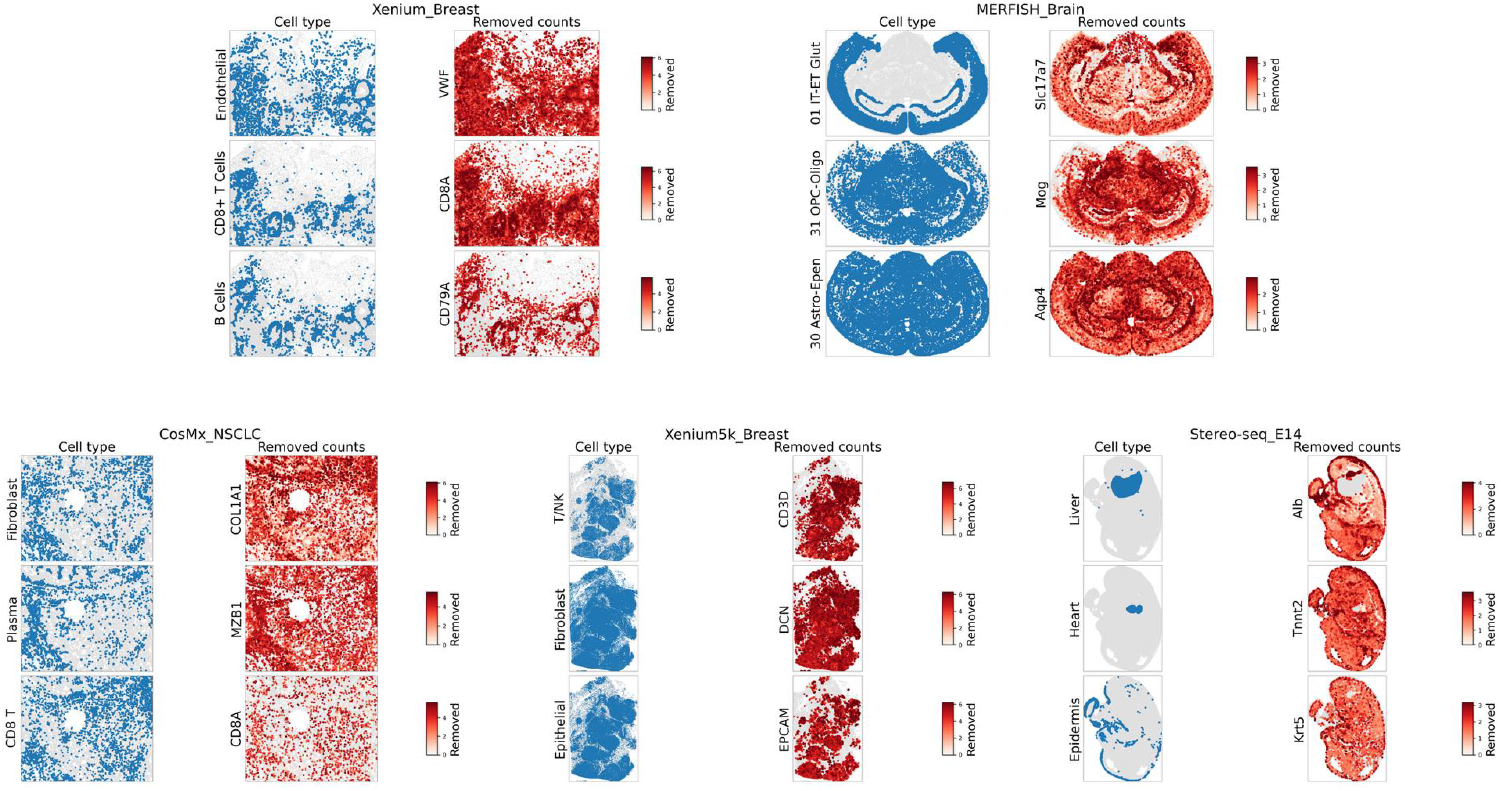
Spatial maps of DeSpotX-decontaminated counts for canonical cell-cluster markers across the five datasets. For each marker, cells of the canonical expressing cluster are highlighted on the left, and the spatial distribution of removed counts is shown on the right.

## M Spatial coherence of baseline methods

We extend the Moran’s *I* comparison in Figure 3 to the four baseline methods (Figure 18). SoupX and DecontX produce scatters that closely follow the identity line, indicating that their decontamination preserves the raw spatial structure of expression but does not enhance it. SpaceBender’s scatters fall on or below the diagonal across all five datasets, suggesting that its decontamination reduces spatial coherence relative to the raw counts. ResolVI shows substantially elevated Moran’s *I* values across all five datasets, often saturating at 0.4–0.8 even for genes with near-zero raw values. This pattern reflects ResolVI’s spatial-smoothing decoder, which incorporates neighbor information into each cell’s expression estimate and inflates Moran’s *I* regardless of whether the underlying signal is biological or ambient. Such smoothing produces visually coherent maps but does not separate native signal from ambient contamination. By contrast, DeSpotX (Figure 3) raises Moran’s *I* above the diagonal proportionally to the raw value, indicating that improvements arise from removing local contamination rather than from spatial averaging.

**Figure 18:**
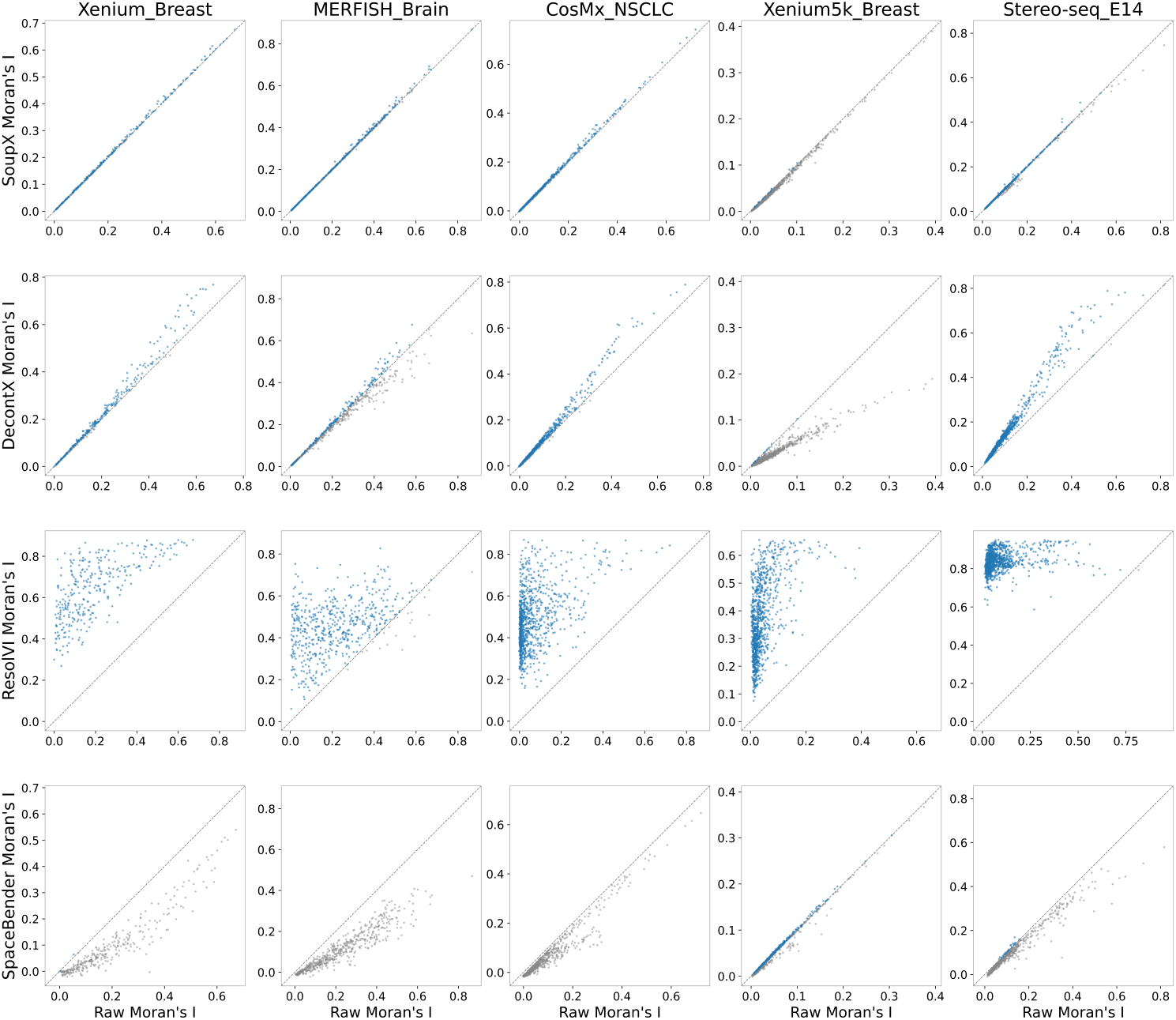
Per-gene Moran’s *I* for raw and decontaminated counts from each baseline method (SoupX, DecontX, ResolVI, SpaceBender) across the five datasets. Each panel plots the raw Moran’s *I* on the x-axis against the decontaminated Moran’s *I* on the y-axis; the dashed line denotes equality.

## N Iterative decontamination on Xenium_Breast

We apply the iterative procedure described in Section 4.3.3 to the `Xenium_Breast` dataset. The CellTypist classifier [22] for this dataset was trained on a published single-cell breast cancer atlas and covers nine broad cell-cluster categories spanning normal and malignant epithelial populations.

The cluster-validity metrics improve significantly from iteration 0 to iteration 1 and stabilize thereafter (Figure 19a), with Silhouette rising from 0.07 to 0.19 and Calinski-Harabasz increasing from 1,590 to 2,089. The ARI between consecutive iterations approaches 1.0 by iteration 2, indicating that the procedure converges to a stable annotation. Additionally, the UMAP embeddings show clearer separation of breast cell types across iterations (Figure 19b).

**Figure 19:**
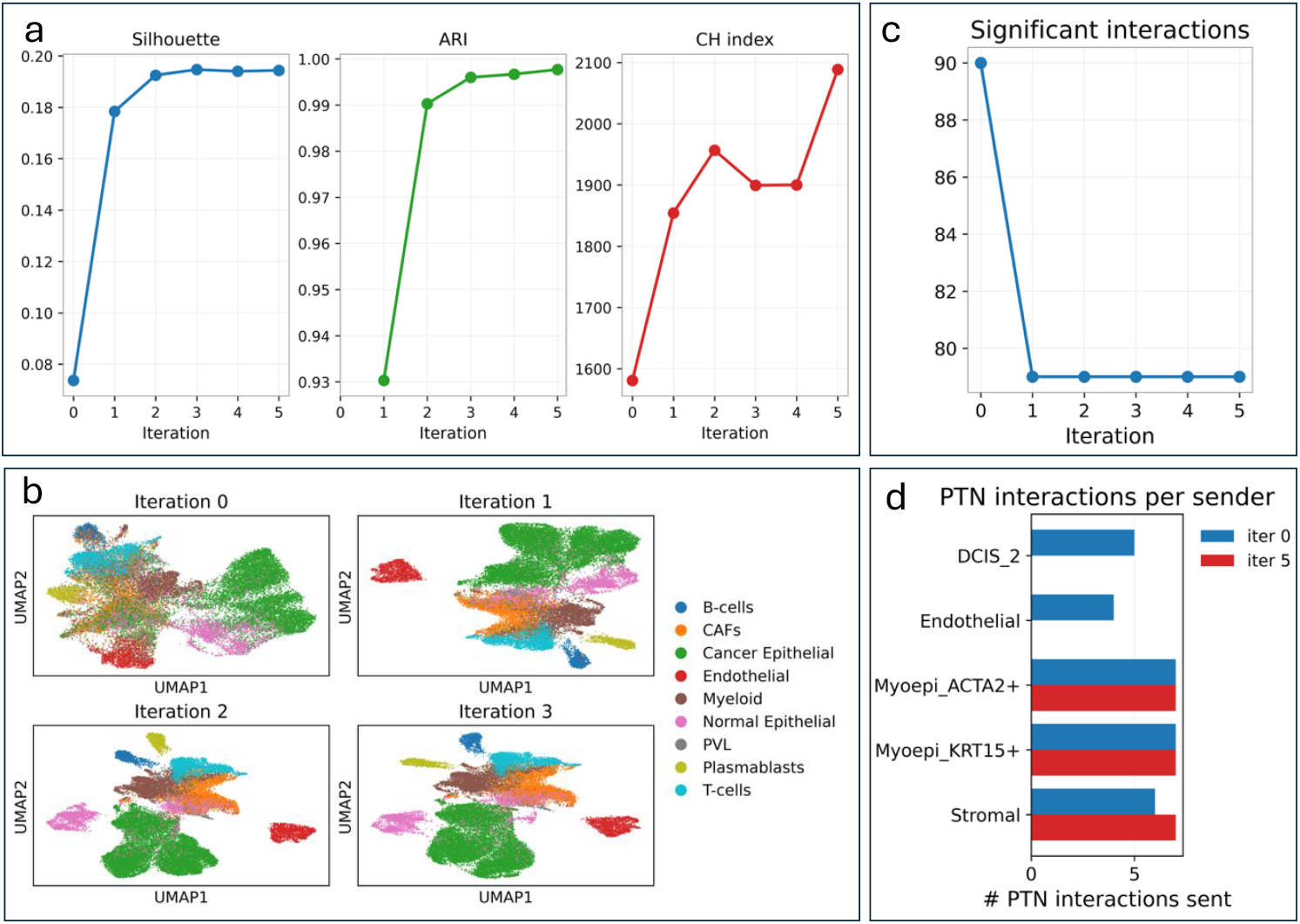
Iterative DeSpotX decontamination and cell-cluster annotation on `Xenium_Breast`. (a) Cluster-validity metrics across iterations. (b) UMAP embeddings of the decontaminated counts at iterations 0-3. (c) Total number of significant cell-cell interactions inferred by CellChat across iterations. (d) Number of significant PTN-pathway interactions per sender cell-cluster at iteration 0 and iteration 5.

CellChat [25] inference shows a reduction in the number of significant interactions between iteration 0 and iteration 1, after which the network stabilizes (Figure 19c). The reduction is largely driven by removal of interactions involving the secreted ligand PTN (pleiotrophin). At iteration 0, PTN is inferred to be sent from cancer (DCIS_2) and endothelial cells; by iteration 5, these interactions are removed, while PTN signaling from myoepithelial cells, which natively express PTN in mammary tissue, is preserved (Figure 19d).

## Notes

### Competing Interest Statement

The authors have declared no competing interest.

